# Genome replication and cell division control by the methylation-sensitive GcrA regulator in *Agrobacterium tumefaciens*

**DOI:** 10.64898/2026.05.29.728657

**Authors:** Florian Fournes, Sandra Martin, Nicolas Pellaton, Michael Taschner, Karolina Bojkowska, Julien Marquis, Stephan Gruber, Justine Collier

## Abstract

Bacteria with complex genomes including secondary chromosomes and/or megaplasmids face unique challenges when controlling their cell cycle: they not only need to coordinate genome replication with other cell cycle events such as cell growth and cell division, but also to synchronize the replication of different replicons. In *Alphaproteobacteria*, replication and maintenance of large extrachromosomal replicons are usually dependent on RepABC modules, as seen for the plant pathogen *Agrobacterium tumefaciens*. In this study, we demonstrate that the conserved GcrA protein is an essential methylation-dependent transcriptional regulator playing at least two critical roles during the *A. tumefaciens* cell cycle. First, GcrA is required for the on-time and synchronized firing of the three RepABC-dependent origins of replication of *A. tumefaciens* that are needed for chromosomal and megaplasmid replication. Second, GcrA triggers the expression of several essential cell division genes for the timely assembly of a functional divisome. These findings highlight how this conserved global regulator can control and then synchronize a variety of essential cell cycle events in *Alphaproteobacteria* with complex genomes, notably coordinating the maintenance of several RepABC-dependent replicons with cell division.

**Graphical abstract:** 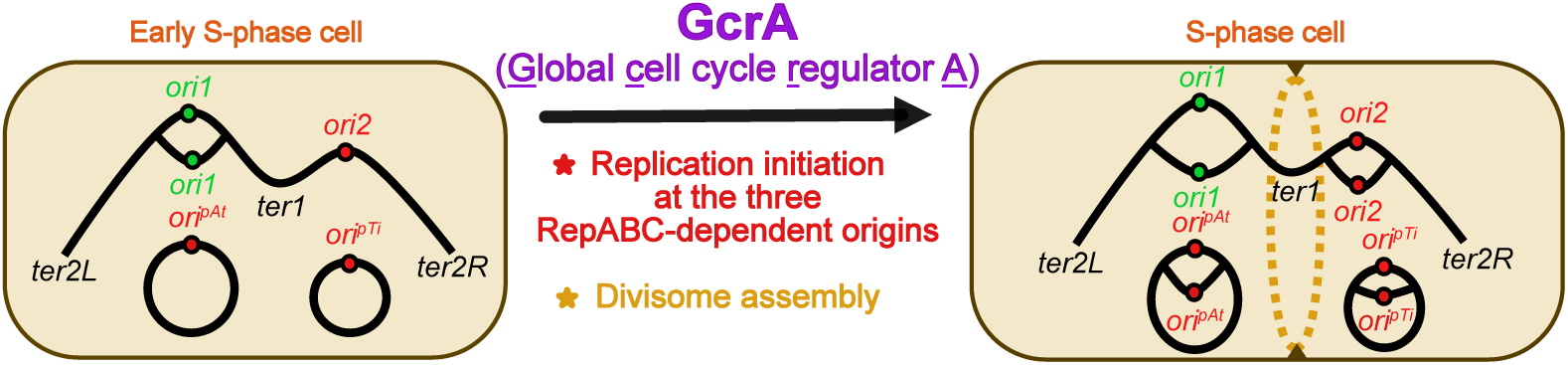

## Introduction

The *Agrobacterium tumefaciens* plant pathogen is a model organism to study *Alphaproteobacteria* with multipartite and/or multicentric genomes (1–4). In such bacteria, representing approximately 10-15% of sequenced bacterial species, including many pathogens or symbionts, core genes are typically distributed across at least two essential chromosomes or on single multicentric chromosomes with multiple replication origins (5,6). These complex genomes also often contain plasmids, including large ones known as megaplasmids, usually involved in environmental adaptation and/or virulence (6–8). Cells with multipartite/multicentric genomes face unique challenges to maintain the integrity of their genome over generations. In these cases, bacteria need to coordinate replication initiation from diverse co-existing origins, while at the same time couple genome duplication with other events of the cell cycle such as growth and cell division.

In all *Alphaproteobacteria* with multipartite or multicentric genomes, the maintenance of each large secondary replicon, such as essential secondary chromosomes or megaplasmids, is governed by a dedicated *repABC* module (9,10). As an example, the genome of the *A. tumefaciens* C58 strain is composed of two essential chromosomes (the primary circular Ch1 and the secondary linear Ch2) together with two non-essential megaplasmids (pAt and pTi) (1,2). Recent studies however revealed that a subset of natural and laboratory-derived *A. tumefaciens* strains carry a single dicentric linear chromosome instead of two distinct chromosomes, likely originating from inter-chromosome recombination (3,4). Importantly, in such strains, the two chromosomal origins located on this unique dicentric chromosome (*ori1* from Ch1 and *ori2* from Ch2) remain essential for viability (3). In *Alphaproteobacteria* with RepABC-dependent replicons, the exact mechanisms coordinating the initiation of replication from the conserved DnaA-dependent *ori1* with initiation from the other RepABC-dependent origin(s) remain elusive, highlighting the need to understand how the replication of RepABC-dependent replicons is temporally controlled during the bacterial cell cycle.

Encoded by *repABC* operons, RepA and RepB bind to *parS* sites (typically located upstream of the *repABC* operon) and are required for replicon partitioning (11,12). As for RepC, it is recruited to a replication origin which is usually embedded within the *repC* coding sequence where it initiates replication (10,13,14). Additionally, certain *repABC* modules were shown to produce a small anti-sense RNA, named RepE, that is transcribed from a region located between the *repB* and *repC* open-reading frames (ORF) (9,15,16). RepE then acts as a negative regulator of replication initiation by repressing the transcription and the translation of *repC* (17,18). Interestingly, previous studies showed that *repABC* modules from megaplasmids or chromosomes are sufficient for replication, but only in certain *Alphaproteobacteria* (12–14,16). This suggests that Alphaproteobacterial actors or regulators may be required for their *in vivo* function. Among these, the <u>c</u>ell <u>c</u>ycle-regulated DNA <u>m</u>ethyltransferase CcrM was recently identified as a potent candidate since CcrM-defective cells display replication defects at the essential RepABC^Ch2^-dependent *ori2* of *A. tumefaciens* (16).

CcrM was first characterized in the environmental *Alphaproteobacterium Caulobacter crescentus* that has one circular chromosome with a single DnaA-dependent origin of replication (19). In this bacterium, CcrM functions only in G2-phase cells towards the very end of the cell cycle, when it methylates adenines (6mA) in 5’-GANTC-3’motifs. During the S-phase of the cell cycle, 5’-GANTC-3’ motifs become hemi-methylated when they get replicated and then remain in that state until the end of the S-phase (20). The presence of 6mA on the *C. crescentus* genome is functionally linked with the activity of the <u>g</u>lobal <u>c</u>ell cycle regulator GcrA that is accumulating during the S-phase of the cell cycle (21). GcrA is a non-canonical transcriptional regulator that binds to methylated 5’-GANTC-3’ motifs located in promoter regions and that interacts with the housekeeping Sigma factor to promote transcription initiation during the S-phase of the *C. crescentus* cell cycle (22–25). Genes activated by GcrA include many involved in cell cycle progression. In particular, the CcrM-GcrA epigenetic tandem plays a central role in the control of cell division, as it promotes the expression of several cell division genes, including *ftsZ,* during the S-phase of the cell cycle (22,23,26,27). FtsZ is a tubulin homolog that polymerizes into a contractile Z-ring at mid-cell, serving as a scaffold for the recruitment of downstream division proteins, such as FtsA, FtsQ and FtsW, forming the divisome (28). Notably, the CcrM/GcrA tandem is conserved across nearly all *Alphaproteobacteria* including those with complex genomes (29–31). In such bacteria, the GcrA regulon has only been described in *Brucella abortus*. In this pathogen, it also preferentially binds to promoter regions with 5’-GANTC-3’ motifs, it is essential for cell division and cell viability and it plays an important role in regulating DNA repair genes (32). Thus, studying the diverse roles of GcrA in different *Alphaproteobacteria* can reveal interesting new epigenetic regulatory pathways (31).

In *A. tumefaciens*, the transcription of the *repABC^Ch2^* operon and the expression of its RepE^Ch2^ anti-sense RNA appeared as being impacted by CcrM-dependent methylation (16), suggesting that GcrA may affect the timing or the frequency of replication initiation at the essential *ori2* although this was never tested directly and the exact mechanisms remained elusive. Strikingly, there is generally a high density of 5’-GANTC-3’ motifs in the promoter regions of *repABC* and *repE* in *repABC* modules (9,16), indicating that the CcrM/GcrA tandem might play a conserved role during the regulation of secondary RepABC-dependent replicons in *Alphaproteobacteria*.

In this study, we shed light on the diversity of the roles of GcrA in *A. tumefaciens* showing that GcrA is essential, not only influencing genome replication, but also other events of the cell cycle such as cell division. We used a combination of transcriptome and live-cell microscopy analyses on a conditional *gcrA* mutant of *A. tumefaciens* to show that: (i) GcrA plays a central role in regulating replication initiation at its three RepABC-dependent origins, and that (ii) GcrA is also critical to control the cell division process. Complementary *in vitro* and transcriptional reporter-based assays further demonstrated that the *A. tumefaciens* GcrA regulator is a methylation-sensitive DNA binding protein that can either activate or repress gene expression. It notably acts as a direct regulator of the expression of *repE^Ch2^*, *repABC^Ch2^* and several essential genes encoding cell division proteins in *Agrobacterium tumefaciens,* so that genome replication and cell division take place at the right moment of the cell cycle. Thus, these findings uncover a novel epigenetic mechanism coordinating genome maintenance with cell division in an *Alphaproteobacterium* with a complex genome.

## Materials and methods

### Bacterial strains and growth conditions

Bacterial strains (*A. tumefaciens*, *C. crescentus* and *Esherichia coli*) used in this study are listed and described in Table S1. Growth conditions are described in the Supplementary Information.

### Plasmids

Plasmids used or constructed in this study are described in Table S2. When applicable, their construction is detailed in the Supplementary Information together with a list of primers used for these constructions (Table S3) and their complete nucleotide sequence is provided as supplementary .dna files when applicable.

### Immunoblotting procedures

Cells pelleted from bacterial cultures (JC2141 and JC2899 strains cultivated in exponential phase in ATGN +/- taurine) were frozen in liquid nitrogen and resuspended in Laemmli buffer (4% SDS, 5% β-mercaptoethanol, 20% glycerol, 0.004% bromophenol blue, 0.125 M Tris-HCl, pH 6) while normalizing samples to the same OD_600nm_. Samples were then boiled at 95°C (10 minutes) before being loaded onto a 12.5% SDS-PAGE polyacrylamide gel. Separated proteins were then transferred onto polyvinylidene difluoride (PVDF) membranes using a PowerBlotter system (Thermo Fisher Scientific). Membranes were then blocked in 5% non-fat dry milk in TBS with 0.05% Tween-20 (TBS-T) and subsequently incubated overnight at 4°C with primary antibodies: custom-made anti-GcrA from rabbits (1:1000; prepared as described in the Supplementary Information) and commercial anti-RpoA from mice (1:7500 from BioLegend, USA). After several washes with TBS-T, membranes were incubated with secondary antibodies: anti-rabbit (1:7,500; Sigma-Aldrich, USA) or anti-mouse (1:7,500; Sigma-Aldrich, USA), respectively. Membranes were washed again three times with TBS-T. Proteins were then detected using the SuperSignalTM West Pico PLUS Chemiluminescent Substrate (Thermo Fisher Scientific, USA) and membranes were imaged using a FUSION FX instrument (Vilber Lourmat, France).

### Microscopy experiments

To analyze the morphology of cells, bacterial cells were first fixed (150 mM NaPO_4_, 12.5% formaldehyde at pH7.5). Fixed cells were then immobilized onto a 0.5× PBS (phosphate-buffered saline) + 1% agarose pad and imaged using a Plan-Apochromat 100×/1.45 phase contrast (Ph3) objective on an AxioImager M1 microscope (Zeiss) with a 1K EMCCD camera (Photometrics) controlled by the VisiView 7.5 software.

For phase contrast combined with fluorescence microscopy analyses, live cells were immobilized and imaged as described above without fixation. Images were then analyzed using the Fiji 2.3.0 software with the MicrobeJ plugin (33) to determine the subcellular localization and the number of fluorescent foci in individual cells. For time lapse imaging (phase contrast and fluorescence microscopy), precultured cells were spotted onto an ATGN +/- taurine agarose (1%) pads. Cells were imaged every 15 minutes using a 100x/1.40 inverted objective on a Leica DMi8 microscope with a sCMOS DFC9000 (Leica) camera and a SOLA light engine (Lumencor).

### RNA extraction for RNA-sequencing (RNA-Seq) and qRT-PCR analyses

RNA samples were prepared from pelleted and frozen cells using the RNeasy Mini-Kit (Qiagen) and adding a DNase I treatment (Qiagen). Resulting RNA samples were treated with a TURBO DNA-free kit (Invitrogen). The quality and the concentration of RNA samples were verified/measured as described in the Supplementary Information. qRT-PCR experiments were performed and analyzed as described in (16). RNA-Seq experiments and analyses are described in the Supplementary Information.

### Promoter activity measurements

The pOT1e derivatives listed in Table S2 were introduced into JC450 or LS3707 *C. crescentus* cells by transformation. The resulting strains were cultured overnight at 28°C in PYE + 0.03 % xylose and diluted into M2G + 0.03 % xylose for another overnight culture. Cultures were then diluted again into M2G to reach an OD_600nm_∼0.05 and transferred into 96-well-plates +/- 0.03 % xylose. Fluorescence intensities and OD_600nm_ were measured after 3 and 6 hours of growth at 28°C using a microplate reader (Biotek Synergy H1: excitation at 479 nm and emission at 520 nm). Relative fluorescence units (RFU) were calculated as the fluorescence intensities (AU)/OD_600nm_. Promoter activities were estimated by calculating the ratio between the RFU of strains carrying pOT1e derivatives with studied promoters and the RFU of the control strain with the empty pOT1e cultivated in the same conditions.

### Fluorescence anisotropy measurements

The procedure used to express GcrA-CDP- His8 in *E. coli* cells and to purify GcrA-CDP-His8 is described in the Supplementary Information. DNA probes were created by annealing two complementary oligonucleotides, of which only one was labelled at its 3’-end with 6-Carboxyfluorescein (6-FAM). Each oligonucleotide was ordered with 6mA or non-modified A in its 5’-GANTC-3’ motifs (see Table S3) to create fully methylated, non-methylated or hemi-methylated DNA probes (all four possible combinations) after the annealing procedure. DNA/protein binding experiments were carried out in an FA binding buffer containing 10 mM Tris-HCl pH 7.5, 100 mM KCl and 100 µg/ml salmon sperm DNA (ThermoFisher). Measurements were recorded using a Synergy Neo Hybrid Multi-Mode Microplate reader (BioTek) equipped with appropriate filters in black 96-well flat bottom plates at 25°C. The volume added to each well was 50 µl. DNA probes were always added at a concentration of 50 nM. The GcrA-CPD-His8 protein was titrated by first creating a condition with high concentration (typically 2.5 - 5 µM) and subsequently creating 2-fold serial dilutions. One well was always kept free of protein to obtain a baseline for the unbound DNA probe. All measurements were carried out in triplicates. Anisotropy values were obtained directly from the BioTek Synergy Neo software, exported, and then fit using non-linear regression in GraphPad Prism.

### Statistics and reproducibility

Statistical methods and sample sizes (n) are described in corresponding figure legends when necessary. Statistical analyses were done using the either Excel, GraphPad-PRISM or R softwares.

## Results

### GcrA depleted cells display early cell division defects and die over time

We previously constructed a conditional *gcrA* mutant derived from the *A. tumefaciens* C58 strain that has one dicentric chromosome and two megaplasmids (16). In this strain, the only copy of *gcrA* was inserted at the *tetRA* locus under the control of the P*tauA* promoter (Δ*gcrA PtauA-gcrA*) so that GcrA can be depleted from cells when cultivated in media without the taurine inducer. Using this strain, we previously found that GcrA-depleted cells remain viable for a few hours when cultivated in minimal medium (ATGN) and accumulate lower than normal levels of the *repC^ch2^*mRNA (16). Beyond this, the precise phenotypes and the transcriptome of GcrA-depleted cells were not characterized. Here, we used this strain to investigate what happens before cells die when GcrA is depleted to shed light on why GcrA is essential. First, GcrA protein levels were measured over time after the removal of the taurine inducer. This revealed a rapid decrease, with GcrA becoming hardly detectable after 2 hours of growth in ATGN medium (Fig. 1A), also demonstrating that GcrA is a relatively unstable protein. To estimate the half-life of GcrA more precisely, shorter time points were also analyzed revealing that it is close to 15 minutes (Fig. S1A) in cells with a doubling time of approximately 154 minutes (Fig. S1B). This short half-life (<15% of the cell cycle duration) is consistent with a potential role of GcrA as a cell cycle regulator in *A. tumefaciens*. Colony-forming unit (CFU) assays then demonstrated that cell viability starts to drop soon after GcrA becomes undetectable (obvious after 6 hours of GcrA depletion in ATGN medium; Fig. 1A&B). Time-lapse microscopy analyses then revealed that GcrA-depleted cells elongate over time (Fig. 1C) and then frequently develop branched cell poles (Fig. 1C&D), which is typical of early cell division defects in *A. tumefaciens* (34). CFU assays (Fig. 1B), corroborated by live/dead fluorescence microscopy (Fig. S2A), revealed that highly branched cells with multiple poles (Fig. S2B) are no longer viable. Notably, very similar phenotypes were observed when GcrA was depleted in cells cultivated in a YEB complex medium, even if higher concentrations of the taurine inducer were required for full complementation (Fig. S3).

**Figure 1:**
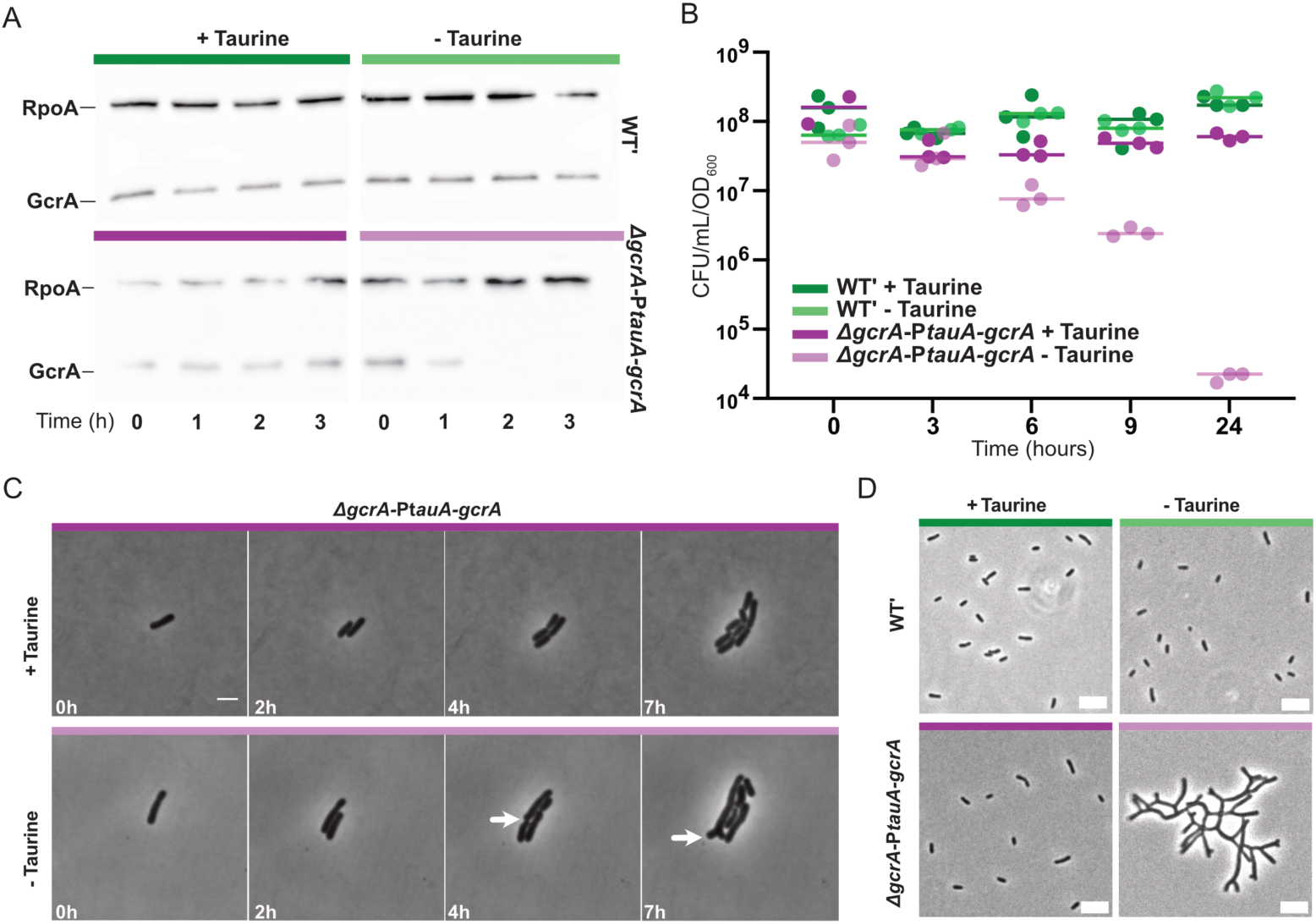
GcrA depletion impacts cell morphology and viability in *A. tumefaciens.* (**A**) Immunoblots showing GcrA levels (and RpoA levels as a loading control) in JC2141 (WT’) and JC2899 (Δ*gcrA PtauA-gcrA*) cells cultivated in ATGN medium +/- taurine for the indicated time after exponentially growing cells (OD_600nm_∼0.3) were pre-cultivated in ATGN + taurine. (**B**) Quantification of colony-forming units (CFU) on ATGNA + taurine from cultures of JC2141 and JC2899 strains in ATGN +/- taurine for the indicated time. Three independent biological replicates were quantified, and each dot represents one replicate; horizontal bars correspond to the mean value of the three replicates. (**C**) Phase-contrast (Ph3) time-lapse microscopy images comparing the morphology of JC2899 cells cultivated in ATGN +/- taurine. Cells were pre-cultivated as in panel B before being dropped onto ATGN +/- taurine agarose pads at time 0. The white arrow shows a cell pole that progressively branches over-time. Scale bars correspond to 2 μm. (**D**) Phase-contrast microscopy images of fixed JC2141 and JC2899 cells cultivated in ATGN +/- taurine for 24 hours. Scale bars correspond to 5 μm. In all panels, taurine was added at 2.5mM when indicated and time is in hours.

Altogether, these results demonstrate that GcrA is essential for the survival of fast- and slow-growing *A. tumefaciens* cells and that cell division is strongly impaired before cell death when GcrA is depleted.

### GcrA is a global transcriptional regulator in A. tumefaciens

GcrA was initially identified as a <u>g</u>lobal <u>c</u>ell cycle regulator functionally linked with CcrM in *C. crescentus* (35). To investigate the width of impact of GcrA on gene expression in *A. tumefaciens*, we compared the transcriptome of JC2899 (Δ*gcrA PtauA-gcrA*) cells cultivated in ATGN +/- taurine for 3 hours (Fig. 2 and Table S4) by RNA-Seq, before cells started losing their viability (Fig. 1B). Upon GcrA depletion, the expression of 486 genes was significantly affected (fold change (FC) > 2 and adjusted *P*-value < 0.01), including 438 downregulated genes (90%) and 48 upregulated genes (10%) (Fig. 2A and Table S4). This first observation indicated that GcrA may be a transcriptional activator in *A. tumefaciens*, as previously suggested in *C. crescentus* where it interacts with the house-keeping Sigma factor to promote transcription initiation at methylated promoter regions (24,25). However, at this point, we could not exclude the possibility that many of these hits were indirect. Among the 486 mis-regulated genes, 36% contain at least one 5’-GANTC-3’ motif (methylated by CcrM) maximum 200 bp upstream of their open reading frame (ORF), a region typically including promoter elements (red dots in Fig. 2A), which is significantly more than randomly expected (24%), suggesting that GcrA may also be a methylation-sensitive regulator in *A. tumefaciens* at least when binding to a subset of these promoters. Still, when comparing the proposed regulon of CcrM (16) with this GcrA regulon (Table S4), we observed an only limited overlap of 11 genes (Fig. S4A) including the *repC^Ch2^* and *repC^pTi^* genes encoding two replication initiators (1,2), the *pleC* gene (*Atu0982*) encoding an important developmental regulator (36) and three genes encoding GGDEF-containing proteins that most likely control the levels of the c-di-GMP effector (37,38) (Fig. S4A). Globally, GcrA has a direct or indirect impact on the expression of a wide variety of genes, including many belonging to COG categories associated with cellular processes and signaling among the GcrA-activated genes and many belonging to the COG categories associated with information storage and processing among the GcrA-repressed genes (Fig. 2B).

**Figure 2:**
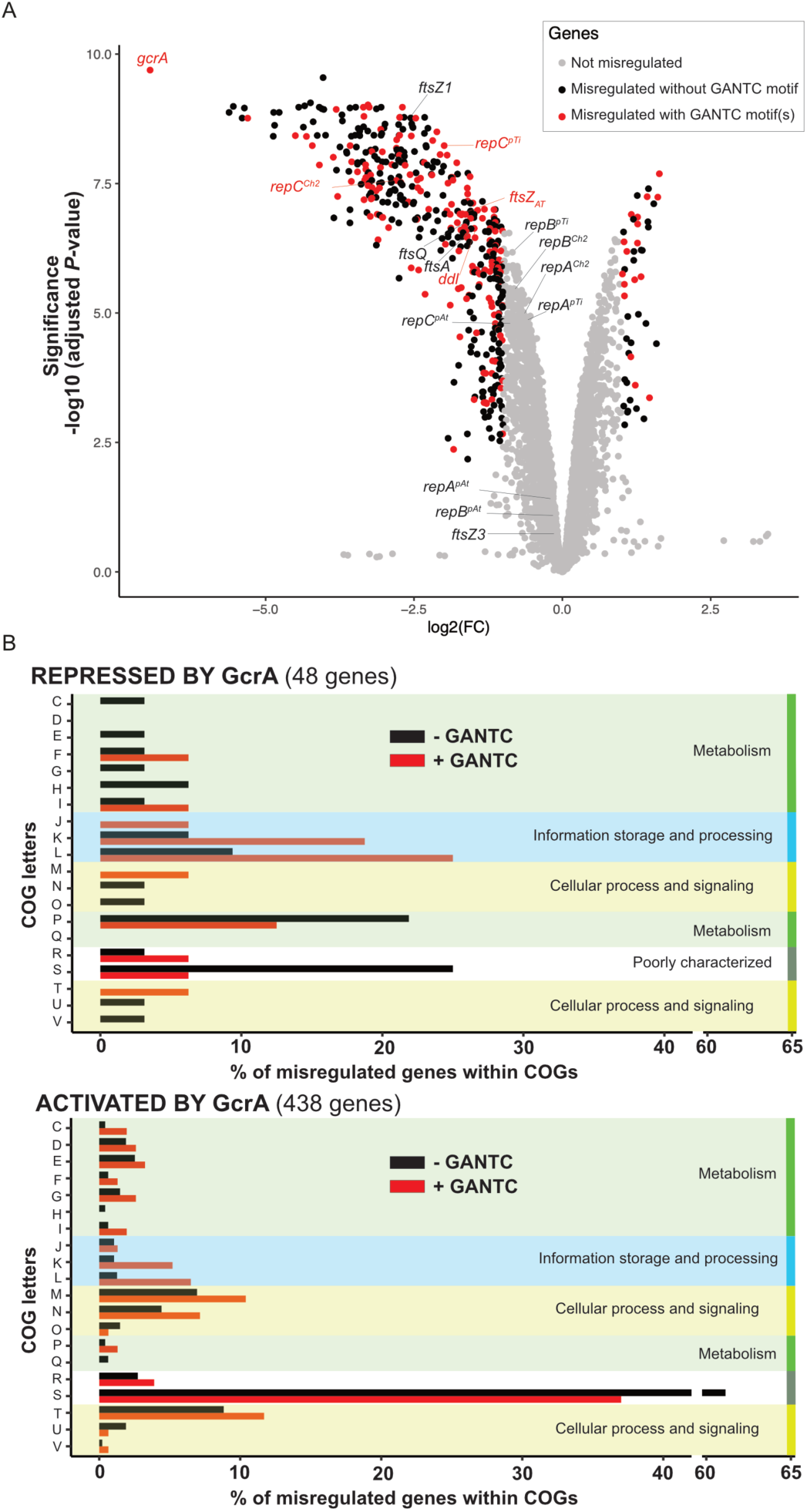
GcrA has a major impact on the *A. tumefaciens* transcriptome. (**A**) Volcano plot comparing the transcriptome of GcrA-depleted and GcrA-repleted cells listed in Table S4. JC2899 (Δ*gcrA PtauA-gcrA*) cells were cultivated exponentially in ATGN +/- taurine for 3 hours as described in Fig. 1A and total RNA were extracted for RNA-Seq analyses. Each colored dot (black, red, grey) corresponds to one gene. Grey dots correspond to genes that are not considered as significantly mis-regulated (FC < 2 or adjusted *P*-value > 0.01). Black dots correspond to genes that are significantly mis-regulated (FC > 2 and adjusted *P*-value < 0.01) but that do not contain a 5’-GANTC-3’ motif within the 200 bp region upstream of the ORF. Red dots correspond to the genes that contain minimum one 5’-GANTC-3’ motif within the 200 bp region upstream of the ORF and that are significantly mis-regulated. Adjusted *P*-values were calculated with three independent biological replicates for each growth condition (+/- taurine). A Glimma Volcano plot (interactive HTML graphic) is available in a Supplementary file. (**B**) Functional classification of genes significantly mis-regulated upon GcrA depletion. Bar charts represent the percentage of genes that are significantly mis-regulated in GcrA-depleted cells within each COG category (y-axis letters with their description in Fig. S4B) (52). The upper panel includes genes repressed by GcrA, while the lower panel includes those activated by GcrA. Red and black bars include genes with and without minimum one 5’-GANTC-3’ motif within the 200 bp region upstream of the ORF, respectively. COG categories were additionally grouped into broader functional classes (indicated on the right).

Altogether, even if the GcrA regulon of *A. tumefaciens* evolved significantly compared to the *C. crescentus* (22,23), *Brevundimonas subvibriodes* (31) and *B. abortus* (32) regulons, it still appears that it acts as a major cell cycle regulator in all *Alphaproteobacteria* where this was investigated. Below, we dissect the specific impact of GcrA on the essential processes of genome maintenance and cell division in *A. tumefaciens* to understand how it functions as a key cell cycle regulator in this bacterium.

### GcrA promotes replication initiation at the three RepABC-dependent origins of A. tumefaciens

The transcriptome analysis described above (Fig. 2A), complemented with previously published qRT-PCR analyses (16), showed that GcrA is a direct or indirect activator of *repC^Ch2^* expression but these did not investigate the impact of GcrA on chromosomal replication initiation by RepC^Ch2^ at *ori2*. To shed light on this hypothesis, we performed fluorescence microscopy experiments to visualize *ori1* and *ori2* localization reporters in Δ*gcrA PtauA-gcrA* cells carrying a *ygfp-parB^pMT1^/parS^pMT1^* system near *ori2* and a *mCherry-parB^P1^/parS^P1^* system near *ori1* (39) and cultivated in ATGN +/- taurine for 3 hours (Fig. S5A). Cells were then sorted as a function of their size and displayed as demographs to see the number and the localization of each origin as a function of cell cycle progression depending on *gcrA* expression (Fig. 3B&C). In wild-type cells, it has been shown that *ori1* and *ori2* co-localize at the old pole of G1-phase cells (Fig. 3A) (16,40). During the G1-to-S phase transition, chromosome replication initiates first at the *ori1* and one of the two newly replicated copies of the *ori1* is rapidly relocated to the new cell pole. A significant while after, replication also starts at *ori2* and one copy of the newly replicated *ori2* is then relocated to the new cell pole, using a mechanism that is supposedly dependent on a physical association between the *ori1*-associated ParB1 and the *ori2*-associated RepB^Ch2^ proteins (39,40). When *gcrA* was expressed from *PtauA-gcrA* (Fig. 3B&C, with taurine), we observed very similar origin firings and choreographies. In contrast, when the taurine inducer of *gcrA* expression was removed, we observed that the duplication/partitioning of the *ori2* could only be detected in significantly longer cells (Fig. 3B, without taurine) compared to *gcrA*-expressing cells (Fig. 3B, with taurine), while the duplication/partitioning of the *ori1* did not appear to be affected (Fig. 3C). This first observation suggested that replication from *ori2* may be delayed in GcrA-depleted S-phase cells. Quantitative analyses of S-phase cells (corresponding to cells with two *ori1* foci as seen in Fig. S5A&B) revealed that GcrA depletion significantly affects replication initiation from *ori2* or *ori2* partitioning: after 3 hours of growth in ATGN without taurine, the proportion of S-phase cells with only one *ori2* focus doubled, while the proportion of S-phase cells with two *ori2* foci dropped approximately two-fold, compared to GcrA-repleted cells (Fig. 3D). Strikingly, when GcrA was depleted for 6 hours, a substantial fraction of the cell population displayed more than two *ori1* foci, while still containing only one *ori2* focus (Fig. S6A&B), indicating that replication from *ori1* proceeded while *ori2* firing/partitioning was still delayed. Altogether, these results demonstrate that the *A. tumefaciens* GcrA protein is essential to promote replication initiation at *ori2* or *ori2* partitioning, while it does not affect initiation at *ori1* or *ori1* partitioning. Interestingly, a significant proportion of GcrA-depleted cells with a single *ori2* focus displayed an unusually positioned *ori2* focus located at non-polar positions, often close to mid-cell, while *ori1* partitioning appeared as normal (Fig. 3B without taurine). It may be that the newly replicated *ori1* that gets segregated at that time of the cell cycle pulls *ori2* away from the old cell pole in GcrA-depleted cells due to the ParB1/RepB^Ch2^ interaction that usually takes place at that moment of the cell cycle.

**Figure 3:**
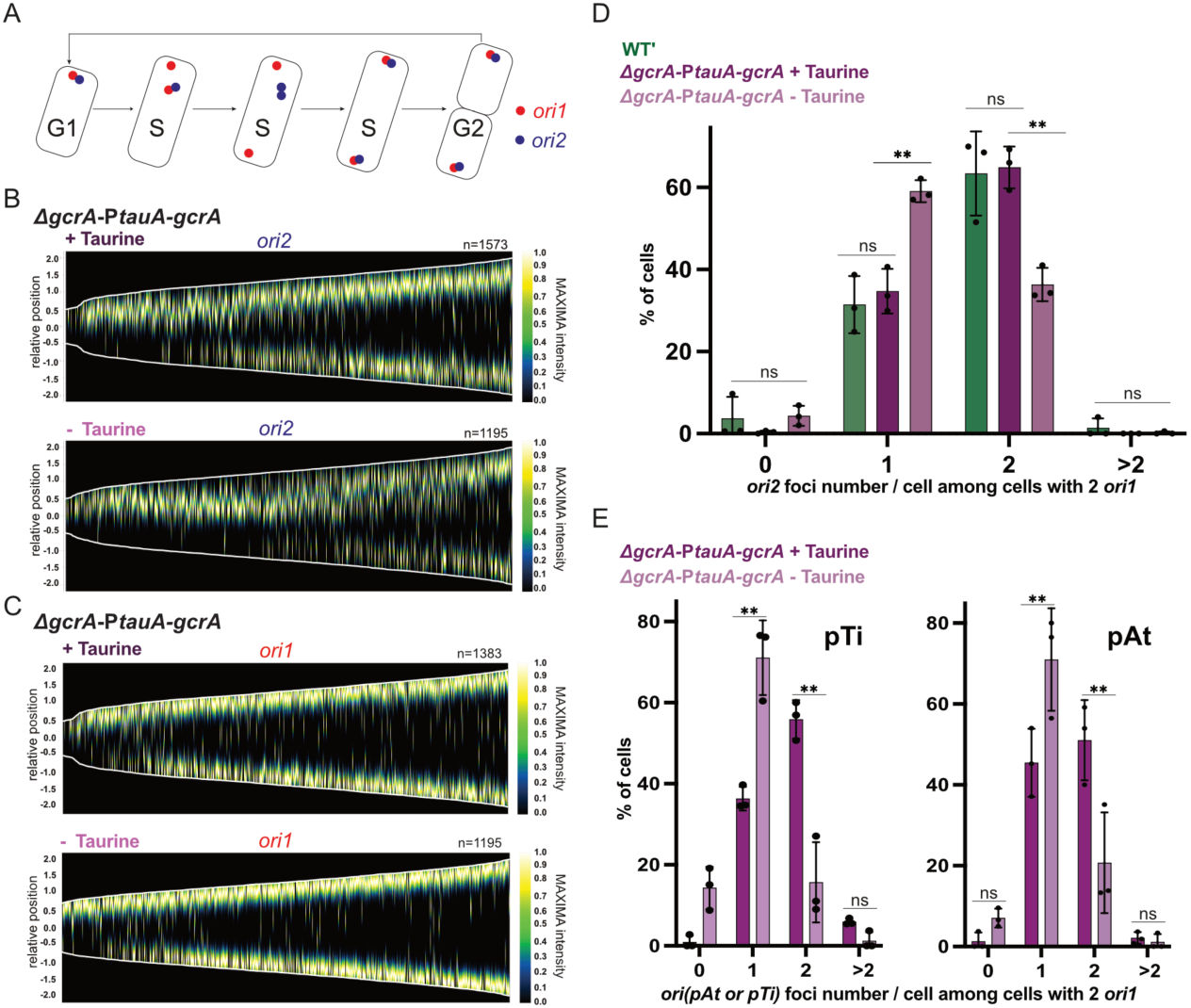
GcrA is critical for chromosome and megaplasmid maintenance in *A. tumefaciens*. (**A**) Schematic describing the intracellular choreography of the replication origins *ori1* and *ori2* during the *A. tumefaciens* cell cycle. (**B)** and (**C**) show demographs obtained from Ph3/RFP/GFP images of JC3095 (JC2899 with *ori1/mcherry* and *ori2/ygfp* reporters) cells cultivated exponentially in ATGN +/- taurine for 3 hours shown in Fig. S5A. Relative position = 0 corresponds to mid-cell; cells measuring from 1-4 μm-long were included into these demographs; n corresponds to the number of cells obtained from three independent replicates and used to construct each demograph. (**D**) Histogram showing the proportion of S-phase cells (with 2 *ori1* foci) that contain a given number of *ori2*. It was obtained using the same data sets as those used in panels (B) and (C), together with data sets obtained with WT’ cells carrying the same reporters (strain JC2777) and cultivated in identical conditions. (**E**) Histograms showing the proportion of S-phase cells (with 2 *ori1* foci) that contain a given number of *ori^pTi^* (left panel) or *ori^pAt^* (right panel). These were obtained using the same data sets as those shown in Fig. S7C&D. JC3096 (JC2899 with *ori1/mcherry* and *oripAt/ygfp* reporters) and JC3097 (JC2899 with *ori1/mcherry* and *oripTi/ygfp* reporters) cells were cultivated and analyzed as described for panel (B), (C) and (D). For histograms shown in (D) and (E): error bars correspond to standard deviations from three independent experiments, Student’s *t*-test: ns = *P*-value > 0.01, ** = *P*-value < 0.01.

As *repC^pTi^* and *repC^pAt^* mRNA levels were also found to be significantly reduced upon GcrA depletion using RNA-Seq (Fig. 2A) and qRT-PCR analyses (Fig. S7A&B), we hypothesized that GcrA may also affect the timing of replication initiation at *ori^pAt^* and *ori^pTi^* in *A. tumefaciens.* The origins of these two megaplasmids usually fire and segregate at the same time as the chromosomal *ori2* (40). As done to visualize the impact of GcrA on initiation at *ori2,* we introduced the *ygfp-parB^pMT1^/parS^pMT1^* reporter near *ori^pAt^* or *ori^pTi^* (40) in Δ*gcrA PtauA-gcrA* cells already carrying the *ori1* fluorescent reporter and proceeded with fluorescence microscopy analyses. These revealed a ∼2-3-fold reduction of S-phase cells containing two *ori^pAt^* or two *ori^pTi^*foci in GcrA-depleted cells (3 hours without taurine) compared to GcrA-repleted cells (Fig. 3E and Fig. S7C&D). Furthermore, nearly 20% of the S-phase cells even appeared to have lost the non-essential pTi megaplasmid (Fig. 3E, left panel without taurine) despite its usually high stability (41).

Collectively, our findings demonstrate that GcrA is essential for replication initiation at the three *repABC*-dependent origins and then that it is a key regulator of genome maintenance in *A. tumefaciens*, which is a novel finding for *Alphaproteobacteria*. This prompted us to investigate how GcrA can promote replication from these three origins at the molecular level.

### GcrA is a methylation-sensitive dual activator of the expression of the RepC^Ch2^ initiator

The transcriptome analysis comparing GcrA-depleted and GcrA-repleted cells revealed that GcrA has a strong positive impact on *repC^Ch2^*mRNA levels (6.6-fold), while it has a more moderate positive impact of *repA^Ch2^*and *repB^Ch2^* mRNA levels (1.6-fold and 1.8-fold, respectively) (Fig. 2A and Table S4). This observation may first appear as surprising since the *repABC^Ch2^* genes are co-transcribed as an operon from a single P_A-Ch2_ promoter upstream of *repA^Ch2^* (16,42) (Fig. 4A). We however demonstrated previously that there is an active anti-sense promoter, named P_E-Ch2_, that is located between *repB^Ch2^* and *repC^Ch2^* (Fig. 4A) that controls the transcription of a sRNA named RepE^Ch2^ and whose activity represses replication from *repABC^Ch2^* (16). A working hypothesis was then that GcrA may not only activate the P_A-Ch2_ promoter, but also repress this P_E-Ch2_ promoter, together leading to a strong activation of *repC^Ch2^* but not *repAB^Ch2^*expression (Fig. 4D).

**Figure 4:**
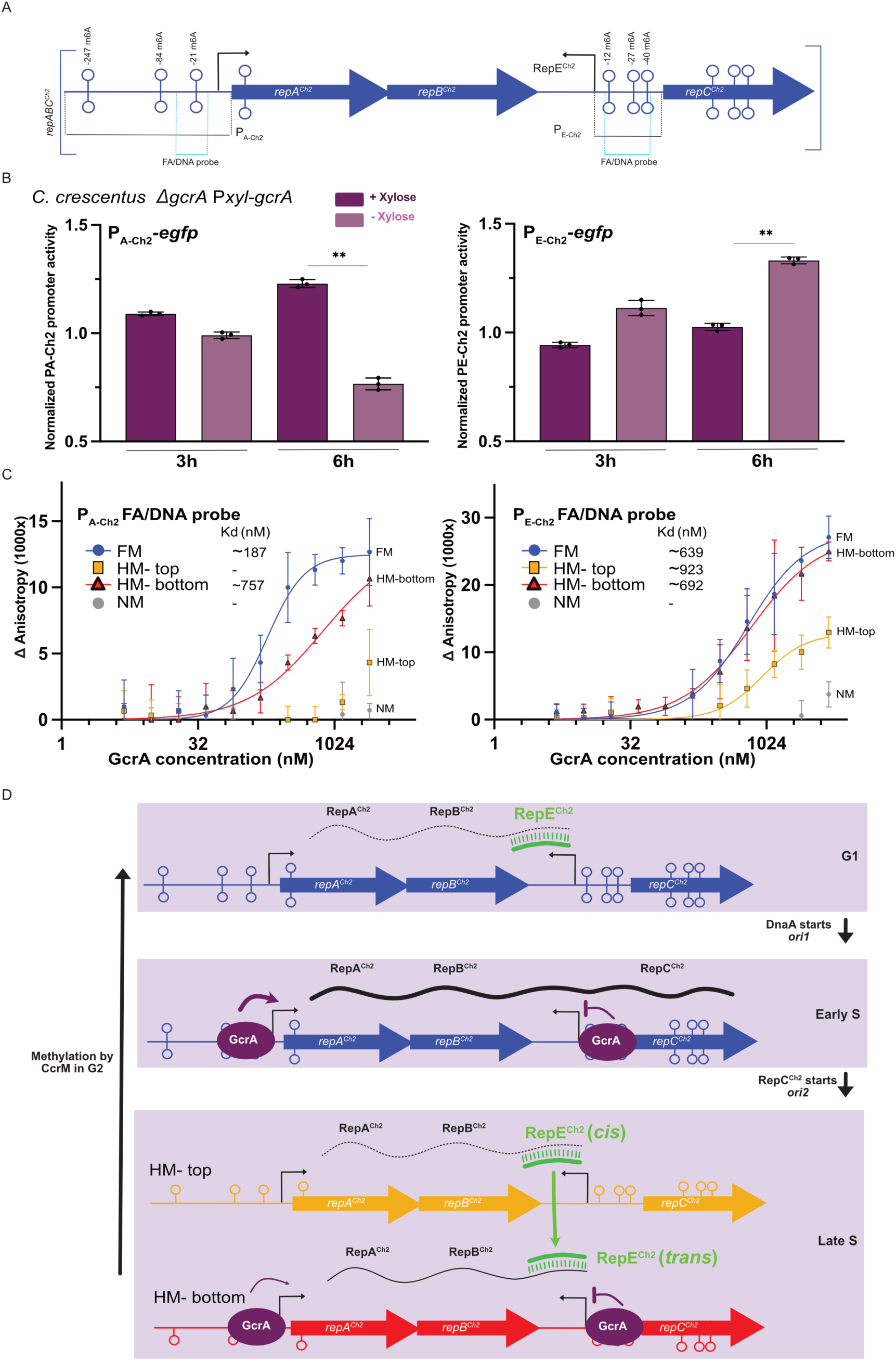
GcrA is a direct methylation-sensitive regulator of *repC^Ch2^* expression. (**A**) Schematic of the *repABC^Ch2^* operon (∼4100 bp) indicating ORFs (blue arrows, not-to-scale) and the 5’-GANTC-3’ motifs (lollipops) that it contains. Black arrows indicate the TSS previously detected in this region in (42). This schematic also shows the P_A-Ch2_ and P_E-Ch2_ promoter regions cloned and used to measure promoter activities in (B) and the P_E-Ch2_ and P_A-Ch2_ FA/DNA probes (light blue lines) designed and used for fluorescence anisotropy (FA) experiments in (C). (**B**) Impact of GcrA_Cc_ on P_E-Ch2_ and P_A-Ch2_ activities in *C. crescentus* cells. The indicated transcriptional reporters (P_E-Ch2_*-egfp* and P_A-Ch2_*-egfp*) were introduced into *ΔgcrA* P*xyl-gcrA C. crescentus* cells (LS3707). Cells were cultivated exponentially in M2G +/- xylose for 3 or 6 hours as indicated. Histograms represent the means of normalized promoter activities (divided by the basal activity detected from the pOT1e empty vector in cells cultivated with xylose) from three independent experiments for each condition; error bars correspond to standard deviations. Statistical significance was assessed for the 6-hour time point (when GcrA_Cc_ is depleted in the absence of xylose) using Student’s *t*-tests: ns = *P*-value > 0.01, ** = *P*-value < 0.01. (**C**) Fluorescence anisotropy assays measuring the binding of GcrA on selected regions of P_E-Ch2_ and P_A-Ch2_. The GcrA protein of *A. tumefaciens* was tagged (GcrA-CDP-His8) and purified from *E. coli* cell extracts (Fig. S11). The indicated concentrations of GcrA-CDP-His8 were incubated with 50 nM of FA/DNA probes. Binding curves are shown for each probe when a Kd could be estimated. FM corresponds to fully-methylated probes, HM corresponds to hemi-methylated probes (top/bottom as illustrated in (D)) and NM corresponds to non-methylated probes; all 5’-GANTC-3’ motifs included in each probe are in the same methylation state. Bars correspond to standard deviations from means obtained from three independent technical replicates. **(D)** Schematic showing the methylation state of the *repABC^Ch2^* operon (fully-methylated DNA in blue; hemi-methylated DNA in red) depending on cell cycle stages in *A. tumefaciens* (16) and a proposed model describing the impact of GcrA on the activities of P_A-Ch2_ and P_E-Ch2_ as a function of cell cycle/methylation stages. The predicted presence of *repAB^Ch2^* or *repABC^Ch2^* mRNA and the RepE anti-sense RNA (that can potentially act in *cis* or *trans*) are also shown at each stage.

To test this hypothesis, we first examined the potential effect of GcrA on the activity of the P_A-Ch2_ and P_E-Ch2_ promoters using P-*gfp* transcriptional reporters (selected promoter regions are shown in Fig. 4A) in a *gcrA* conditional mutant of *C. crescentus* (*ΔgcrA* P*xyl-gcrA*) (21); we chose to use this heterologous assay in *C. crescentus* instead of *A. tumefaciens* since GcrA is neither essential for viability in M2G medium, nor required for replication control in this *Alphaproteobacterium*, avoiding too many pleiotropic effects (26,27). When GcrA_Cc_ was depleted (mutant cells cultivated in M2G without the xylose inducer of P*xyl*), we observed that P_A-Ch2_ activity decreased over time (>50% lower after 6 hours of GcrA_Cc_ depletion in Fig. 4B, left panel), showing that GcrA activates *repABC^Ch2^* expression. Concerning P_E-Ch2_ activity, we observed the opposite, with an increased P_E-Ch2_ activity over time (∼35% after 6 hours of GcrA_Cc_ depletion in Fig. 4B, right panel), indicating that GcrA_Cc_ can repress the expression of the RepE^Ch2^ anti-sense RNA of the *repABC_Ch2_*operon of *A. tumefaciens*. Collectively, these results indicate that GcrA_At_ most likely has a dual positive impact on *repC^Ch2^* expression through an activation of the P_A-Ch2_ and a repression of the P_E-Ch2_ promoters in *A. tumefaciens*; these impacts of GcrA could still be direct or indirect.

To test if the *A. tumefaciens* GcrA protein is a direct regulator of the P_A-Ch2_ and/or P_E-Ch2_ promoters, we then switched to *in vitro* assays. We purified a tagged version of the GcrA protein of *A. tumefaciens* and then conducted DNA binding assays by fluorescence anisotropy (43) using fluorescein-labelled DNA probes that included selected zones of the core P_A-Ch2_ or P_E-Ch2_ promoters that contained 5’-GANTC-3’ motifs that could potentially be recognized by GcrA (Fig. 4A and their sequence/labeling is shown in Fig. S8). Given that GcrA_Cc_ was shown to be a methylation-sensitive transcriptional regulator in *C. crescentus* (22,23), we used non-methylated (NM), hemi-methylated (HM) and fully-methylated (FM) DNA probes with or without 6mA at each 5’-GANTC-3’ motif. Indeed, we found that binding to these promoter probes was strictly methylation-dependent. Specifically, we observed that the *A. tumefaciens* GcrA protein could bind to the fully-methylated P_A-Ch2_ promoter probe with 6mA on both strands of its unique regulatory “M3” 5’-GANTC-3’ motif (identified in (16)), but not to the same non-methylated probe (Fig. 4C). Concerning the P_E-Ch2_ promoter, we also observed that GcrA required 6mA bases to bind to the selected P_E-Ch2_ promoter probe that contained three 5’-GANTC-3’ motifs (Fig. 4C).

Altogether, these observations indicate for the first time that GcrA is a methylation-sensitive transcriptional regulator in *A. tumefaciens*, acting as a direct activator of *repABC^Ch2^* transcription and as a direct repressor of RepE^Ch2^ expression at the beginning of the S-phase of the cell cycle (before *ori2* starts replication), when the *repABC^Ch2^* locus is still in a fully-methylated state (Fig. 4D). Considering that RepE may not only repress *repC^Ch2^* transcription, but also *repC^Ch2^* translation (12), we then predict that GcrA could play a major role in boosting RepC^Ch2^ levels for replication initiation at *ori2* at that specific time of the cell cycle.

### GcrA activates cell division genes and is required for divisome assembly in A. tumefaciens

To faithfully transmit their genetic information, bacteria need to coordinate genome replication with cell growth and cell division. Considering that GcrA appears to regulate several replication initiators and important cell division genes (Fig. 2A, Table S4), we hypothesized that GcrA may play a crucial role in coordinating these essential events of the cell cycle. Consistently, GcrA-depleted cells not only displayed genome maintenance defects (Fig. 3), but also early cell division defects (Fig. 1C&D).

Considering that the *ftsZ_AT_* gene (*Atu2086*), encoding the first divisome component assembled at the septum, was found to be strongly down-regulated (3.2-fold) in GcrA-depleted cells (Fig. 2A), we further investigated divisome assembly. To do so, we leveraged the presence of three different FtsZ homologs in *A. tumefaciens*: FtsZ_AT_ that is essential for cell division and recruits the divisome, FtsZ1 that localizes at the septum in a FtsZ_AT_-dependent manner, and FtsZ3 that is neither required for cell division nor assembled at the divisome (34). To test if FtsZ_AT_ levels become too limiting for divisome assembly in GcrA-depleted cells, we introduced a plasmid carrying a *ftsZ1-sfgfp* fusion under the control of the constitutive *Pvan* promoter (34) into the wild-type and GcrA-depletion strains, enabling us to visualize and compare divisome assembly by fluorescence microscopy. These experiments showed that the early assembly of FtsZ1 at the divisome is significantly impaired when GcrA is depleted for 3 hours in ATGN (Fig. 5A with same conditions as during the RNA-Seq experiments in Fig. 2A). Time-lapse microscopy experiments further showed that even in cells that could still form FtsZ1-sfGFP foci, these foci often faded away over time and/or mis-localized towards a cell pole, where cell branching was then often observed after 5-6 hours of GcrA depletion instead of dividing at mid-cell (Fig. 5B). Furthermore, when looking more carefully at those cells that did display a detectable FtsZ1-sfGFP focus during time-course experiments, we observed that foci where frequently mis-localized along the cell axis, instead of clustering towards mid-cell (Fig. 5C). These observations are strikingly similar to what happens during FtsZ_AT_ depletion (34), indicating that FtsZ_AT_ levels do become limiting for cell division in GcrA-depleted cells.

**Figure 5:**
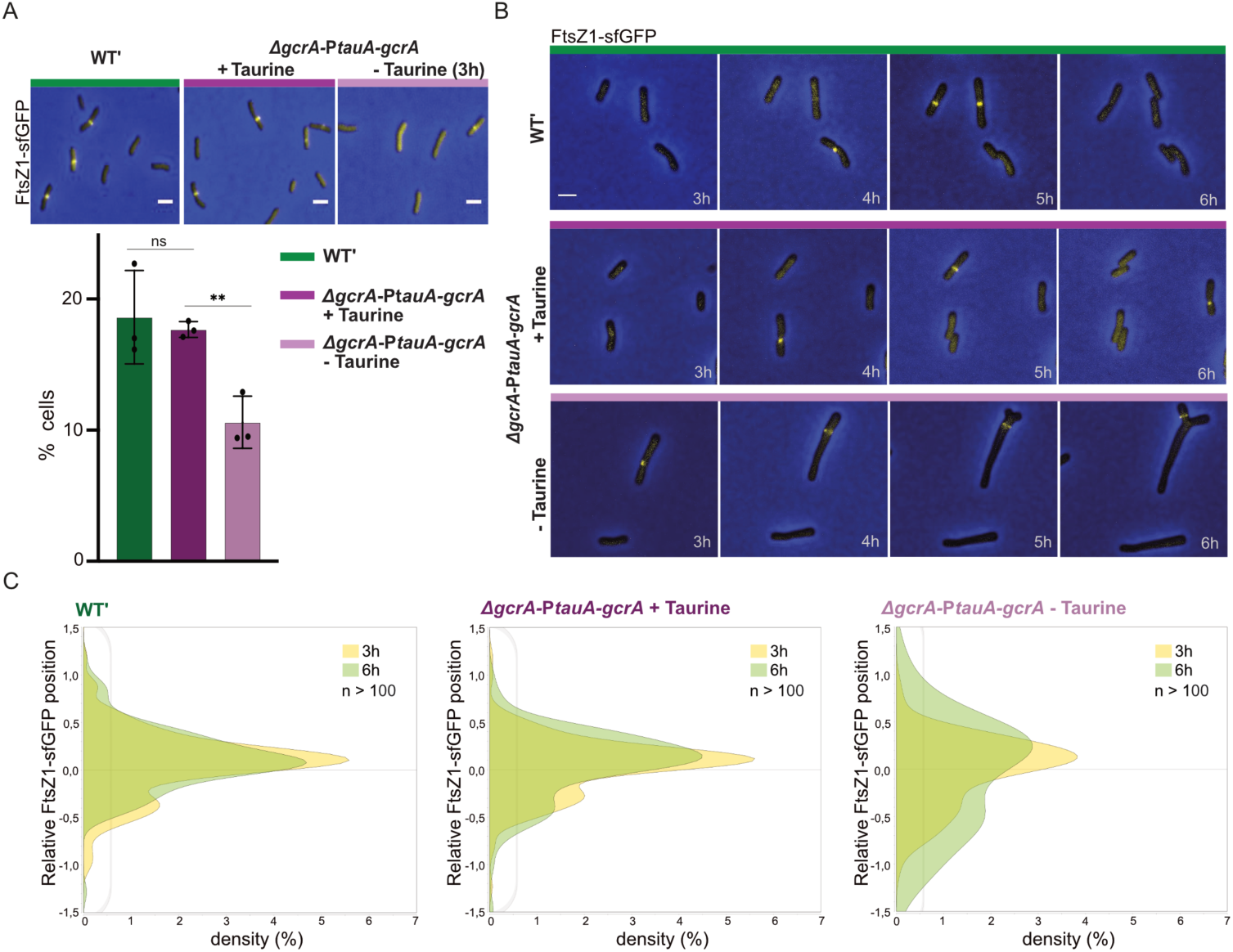
GcrA is required for the early assembly of FtsZ1-sfGFP at the *A. tumefaciens* divisome. (**A**) Divisome assembly is defectious in GcrA-depleted cells. The pRV-P*van*-*ftsZ1-sfgfp* plasmid was introduced into JC2141 (WT′) and JC2899 (Δ*gcrA PtauA-gcrA*) cells and cells were cultured into ATGN +/- taurine for 3 hours. Upper panel: overlap of representative microscopy images (Ph3/GFP). Lower panel: histogram showing the percentage of cells with a FtsZ1-sfGFP focus among 3–5 μm-long cells in the population (pre-divisional or G2 cells from populations as shown in upper panel). Each dot corresponds to an independent replicate and error bars correspond to standard deviations; statistical significance was assessed using Student’s *t*-tests: ns = *P*-value > 0.01, ** = *P*-value < 0.01. (**B**) The divisome is often unstable in GcrA-depleted cells. Time-lapse fluorescence microscopy images of cells cultivated as described in (A) and then transferred onto ATGN +/- taurine agarose pads for 3 more hours. (**C**) The divisome is often mis-localized in GcrA-depleted cells. Fluorescence microscopy images as shown in (A) and Fig. S9 were analyzed to compare the sub-cellular positioning (along the cell’s longitudinal axis) of FtsZ1-sfGFP foci (n > 100 for each condition) in cells expressing or not *gcrA* for 3 (yellow) or 6 hours (green). Relative position: 0 corresponds to mid-cell, 1.5 or -1.5 correspond to cell poles. Scale bars correspond to 2 μm.

Looking more carefully at the *ftsZ_AT_* chromosomal region on the *A. tumefaciens* chromosome, we noticed that *ftsZ_AT_*may belong to the same operon as the four other cell division and cell wall biogenesis genes *ddl*, *ftsQ*, and *ftsA* (Fig. 6A) since it is also the case in several *Bartonellaceae* (44). In line with this, we also noticed that all four genes were similarly activated by GcrA during the transcriptome analyses (Fig. 2A and Table S4: 2.8-fold change, 2.9-fold change, 3-fold change and 3.2-fold change for *ddl*, *ftsQ*, *ftsA* and *ftsZ_AT_,* respectively). To confirm that these genes are co-transcribed, we used cDNA synthesized from whole RNA extracts from wild-type cells to perform PCR analyses (Fig. S10). These demonstrated that *ddl*, *ftsQ*, and *ftsA* and *ftsZ_AT_*belong to the same operon (Fig. 6A). In keeping with this finding, we fused the complete intergenic region between *ftsA* and *ftsZ_AT_* (93 bp, named P_ftsZ*AT*_ in Fig. 6A)), and between *Atu2090* and *ddl* (543 bp, named P_ddl-WT_ in Fig. 6A), with a *gfp* transcriptional reporter and introduced these reporters in *C. crescentus*; we found that P_ddl-WT_ was the only functional promoter region (Fig. 6B). Interestingly, this P_ddl-WT_ promoter region contains five 5’-GANTC-3’ motifs methylated by CcrM and potentially recognized by the GcrA epigenetic regulator to activate this operon (Fig. 6A). To determine which 5’-GANTC-3’ motif(s) may be required for the activation of the P_ddl_ promoter by GcrA in *A. tumefaciens*, we constructed a series of truncated P_ddl_ promoter regions eliminating one 5’-GANTC-3’ motif at a time from the 5’ end of the initial P_ddl-WT_ (Fig. 6A) and measured their activity in the *C. crescentus* conditional *gcrA* mutant. This analysis indicates that the third 5’-GANTC-3’ motif (named M3 and colored in red in Fig. 6A and present in P_ddl-Δ2_, but absent in P_ddl-Δ3_) located ∼84 bp upstream of the *ddl* transcription start site (TSS from (42)) may be required for the activation of the *ddl-ftsAQZ_AT_* operon by GcrA in *A. tumefaciens* (Fig. 6B). To test the potential impact of this M3 on the activation of P_ddl_ by GcrA, we mutagenized this 5’-G<u>A</u>ATC-3’ motif into a non-methylatable 5’-G<u>C</u>ATC-3’motif in the GcrA-activated P_ddl-Δ1_ promoter. Comparing these two constructs, we concluded that this methylatable adenine is probably important for the activation of this cell division operon by GcrA.

**Figure 6:**
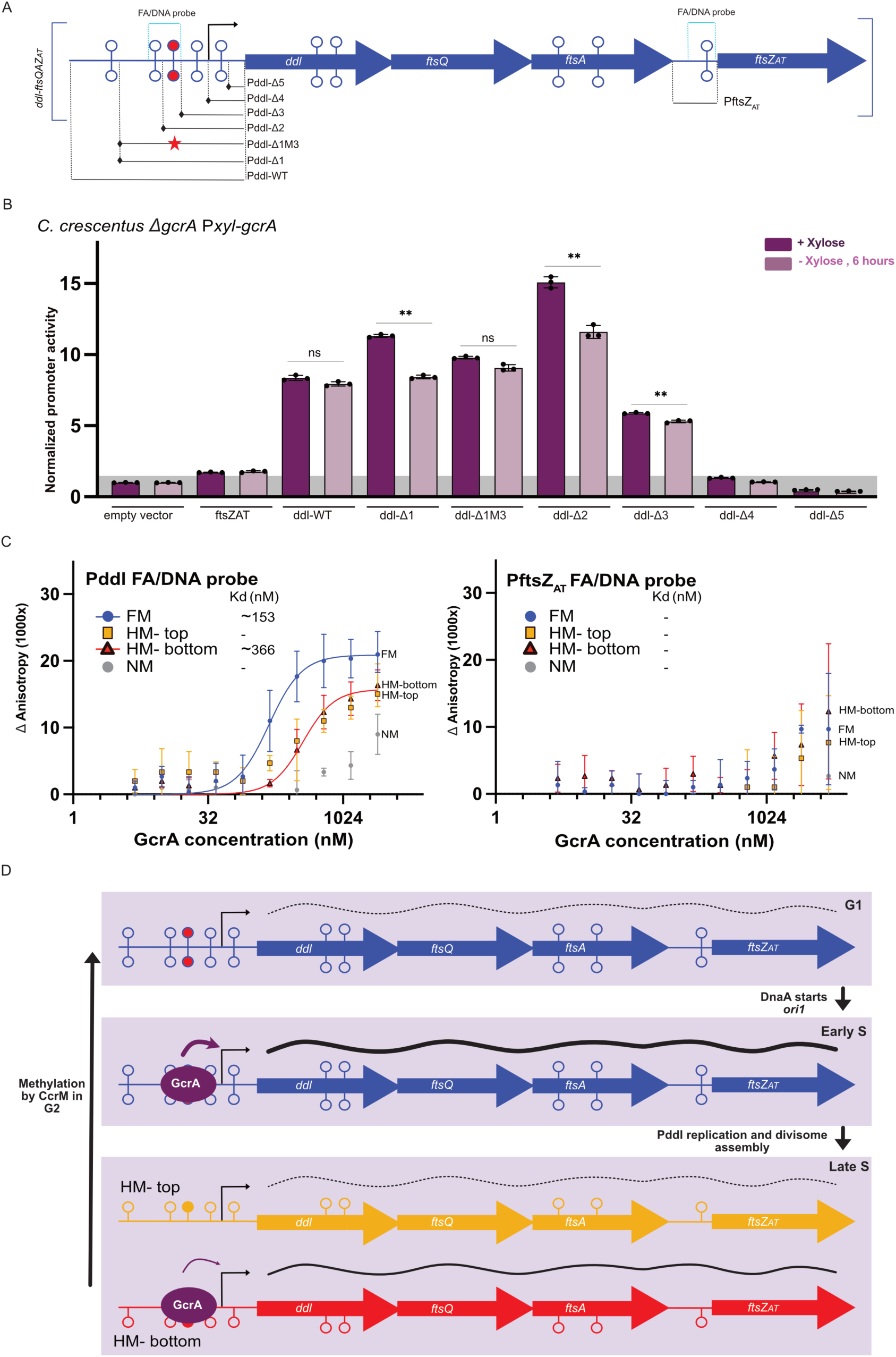
Specific elements of the promoter driving *ddl-ftsQAZ_AT_* transcription are required for activation by GcrA. (**A**) Schematic of the *ddl-ftsQAZ_AT_*operon. ORFs (blue arrows, not-to-scale), 5’-GANTC-3’ motifs (lollipops) and TSS (black arrows from (42)) are indicated. It also shows the cloned P_ddl_ variants and the putative P_ftsZAT_ cloned and used to measure promoter activities in (B) and the FA/DNA probes (light blue) designed and used for fluorescence anisotropy experiments in (C). (**B**) Activities of P_ddl_ variants and of the putative P_ftsZAT_ in *C. crescentus* cells. The indicated transcriptional reporters were introduced into *ΔgcrA* P*xyl-gcrA C. crescentus* cells (LS3707). Cells were cultivated exponentially in M2G +/- xylose for 6 hours (GcrA_Cc_ is depleted in the absence of xylose). Histograms represent the means of normalized promoter activities (divided by the basal activity detected from the pOT1e empty vector in cells cultivated with xylose) from three independent experiments for each construct/condition; error bars correspond to standard deviations. Statistical significance comparing +/- GcrA was assessed using Student’s *t*-tests (only for promoters with an activity above the baseline activity indicated in grey): ns = *P*-value > 0.01, ** = *P*-value < 0.01. (**C**) Fluorescence anisotropy assays measuring the binding of GcrA-CDP-His8 to DNA probes located around the M3 5’-GANTC-3’ motif of P_ddl_ and towards the putative P_ftsZ_ (as illustrated in (A)). Experiments were performed and results were analyzed exactly as described in Fig. 4C except that different FA/DNA probes were used. **(D)** Schematic showing the methylation state of the *ddl-ftsQAZ_AT_*operon (fully-methylated DNA in blue; hemi-methylated DNA in red) depending on cell cycle stages in *A. tumefaciens* (16) and a proposed model describing the impact of GcrA on the activities of P_ddl_ as a function of cell cycle/methylation stages.

We therefore focused on that selected region of the *ddl* promoter to perform fluorescence anisotropy binding experiments using the purified GcrA protein of *A. tumefaciens.* Using differentially methylated P*_ddl_*probes (shown in Fig. 6A, while the sequence/labeling is shown in Fig. S8), we again detected the strongest binding of GcrA onto the fully-methylated (FM) P*_ddl_* probe and the poorest binding to the non-methylated (NM) probe (Fig. 6C, left panel). This observation suggests that GcrA activates the transcription of this essential cell division operon in early S-phase cells when this operon is still fully-methylated (Fig. 6D) and when cells get ready to assemble their divisome at mid-cell (34). As a control, we also tested if GcrA could bind to a DNA probe containing the unique 5’-GANTC-3’ motif found in the IG region upstream of *ftsZ_AT_* (selected region shown in Fig. 6A and the sequence/labeling of the probe is shown in Fig. S8). This assay showed that GcrA could not bind to this IG region, even when it is methylated (Fig. 6C, right panel); this observation shows the high specificity of GcrA for specific regulatory 5’-GANTC-3’ motifs in active promoter regions and not just for random 5’-GANTC-3’ motifs on the *A. tumefaciens* genome.

## Discussion

The findings described in this study demonstrate that GcrA is an essential <u>g</u>lobal <u>c</u>ell cycle regulator in *A. tumefaciens* (Fig. 1&2, Table S4 and Fig. S2&S3) that can sense epigenetic signals (6mA) added by CcrM onto the *A. tumefaciens* genome (Fig. 4&6). 5’-GANTC-3’ motifs methylated by CcrM most likely switch from a fully-methylated state in G1-phase cells, to a hemi-methylated state once they get replicated during the S-phase of the cell cycle; they then stay in this hemi-methylated state for a significant time period until cells reach the G2-phase of the cell cycle when CcrM methylates newly synthesized DNA strands (16). Then, the GcrA/CcrM couple can potentially be used to control the timing of gene expression during the cell cycle and to synchronize gene activation or inhibition with the entry of cells into S- or G2-phases. Here, our data suggests that this property of the GcrA/CcrM couple is exploited by *A. tumefaciens* to dictate when three of the origins of replication of the dicentric chromosome and of its two megaplasmids start replication and to promote the assembly of the divisome at the beginning of the S-phase of the cell cycle so that predivisional cells are then ready to divide on-time.

### GcrA, an epigenetic regulator that can activate or repress the transcription of its target genes in *A. tumefaciens*

Our data shows that the *A. tumefaciens* GcrA protein can directly activate or repress the promoters that it targets, as exemplified with the Pddl/P_A-Ch2_ (Fig. 4&6) and the P_E-Ch2_ (Fig. 4) promoters, respectively. This is an original finding since previous structural data showed that the *C. crescentus* GcrA protein acts exclusively as a transcriptional activator in this bacterium, where it binds to the RNAP/Sigma factor to promote their loading onto target promoters to initiate transcription (24,25).

Beyond this first difference, we also found that the regulon of GcrA in *A. tumefaciens* evolved significantly compared to the regulons of GcrA in *C. crescentus* (21–23) or *B. subvibriodes* (31), and that GcrA is essential for the viability of *A. tumefaciens* not only in fast (in YEB medium in Fig. S3), but also in slow (in ATGN medium in Fig. 1 and Fig. S2A) growing conditions. This essentiality, which is not seen in *C. crescentus* (26) or *B. subvibriodes* (31), may be connected with direct regulation of *repE^Ch2^*and *repABC^Ch2^* expression by GcrA in *A. tumefaciens* (Fig. 4), boosting RepC^Ch2^ levels to initiate DNA replication at its second essential chromosomal origin (Fig. 3).

Another relevant finding is that even if the conserved and essential *ftsZ_AT_* cell division gene of *A. tumefaciens* (34) is part of a complex *ddl*-*ftsQAZ_AT_* operon (Fig. S10) instead of being transcribed into a GcrA-activated monocistronic transcript as in *C. crescentus* (26,45), its transcription is still directly activated by GcrA and in a manner that is still dependent on the methylation state of its distant Pddl promoter. This illustrates the plasticity of regulatory pathways that can adapt to different gene positioning/clustering architectures while still maintaining similar logics and epigenetic mechanisms of regulation. Considering that the Pddl promoter is located ∼0.8Mbp away from the *ori1* of the dicentric chromosome of *A. tumefaciens*, we anticipate that it should switch from a fully-methylated (FM) state to hemi-methylated (HM) states relatively early after the onset of the S-phase of the cell cycle (Fig. 6D), at a time when the FtsZ_AT_ ring should already be assembled at the septum (34) and when new molecules of FtsZ_AT_ are most likely no more required for cell division to proceed. An intriguing observation that we made during our *in vitro* binding assays was that one of the two newly-replicated hemi-methylated Pddl promoters apparently kept some affinity for GcrA (the “HM-bottom” probe in Fig. 6C&D), while the other showed only poor or no binding (“HM-top” probe in Fig. 6C&D). This potential methylation strand specificity opens the possibility of an asymmetric mechanism of regulation where one of the daughter cells may then be more prone to restarting cell division immediately than the other one; it is then tempting to speculate that this might be connected with the asymmetric nature of the *A. tumefaciens* cell cycle giving rise to daughter cells with different fates (one in G1-phase and the other one directly in S-phase and getting ready to divide sooner) (46,47).

### Do GcrA levels fluctuate during the *A. tumefaciens* cell cycle?

In *C. crescentus*, the levels of GcrA are highly regulated at the transcriptional level and by proteolysis so that GcrA accumulates the most during the S-phase of the cell cycle. Our observation that the *A. tumefaciens* GcrA protein is relatively unstable (Fig. 1A and Fig. S1) fits with the concept that GcrA levels could also potentially fluctuate during the *A. tumefaciens* cell cycle. Unfortunately, despite several attempts, the cell cycle of the *A. tumefaciens* C58 strain that we used in this study could not be easily synchronized to determine if GcrA levels fluctuate (data not shown); future studies should for sure address this interesting question. If GcrA is also predominantly found in *A. tumefaciens* S-phase cells, it could add an additional layer of regulation of its target promoters to fine-tune the exact cell cycle period when specific genes (and biological processes) are activated or repressed by GcrA, beyond the already clear impact of the cell cycle-dependent methylation state of at least a subset of its target promoters.

### The GcrA/CcrM couple coordinates the initiation of replication from different origins on the complex *A. tumefaciens* genome

Our findings show that GcrA is not only necessary for the on-time firing of the essential secondary chromosomal origin (*ori2*) relatively soon after the firing of the *ori1* (Fig. 3B&C&D), but also for the simultaneous firing of the two non-essential origins of replication of the pAt and pTi megaplamids (Fig. 3E). This co-regulation by GcrA probably explains why these three origins initiate replication and segregate essentially at the same time of the cell cycle as previously observed (40). For the chromosomal *repABC^Ch2^* module, it looks like GcrA has a dual positive impact on *repC^Ch2^* expression through a direct activation of *repABC^Ch2^* transcription and through an inhibition of the RepE^Ch2^ repressor of *repC^Ch2^* expression (Fig. 4).

Similarly to the Pddl promoter region, our *in vitro* binding assays also suggest that GcrA has more affinity for the “HM-bottom” probes of the P_A-Ch2_ and P_E-Ch2_ promoters than for its “HM-top” ones even if the strongest affinity was by far found for the FM P_A-Ch2_ probe (Fig. 4C&D), suggesting a potentially asymmetric expression of *repC^Ch2^* between the two nascent daughter cells that may boost an earlier initiation at *ori2* in one of the two daughter cells. The globally lower affinity of GcrA for the HM P_A-Ch2_ probes than for the FM P_A-Ch2_ probe (Fig. 4C&D) may also play a role in limiting RepC^Ch2^ over-accumulation during the second half of the S-phase of the cell cycle (after initiation at *ori2*) when it is no longer required. Future studies should focus on addressing whether these assumptions may be true. For the *repABC* modules of the two megaplasmids, GcrA apparently mostly affects *repC^pAt^/repC^pTi^* expression through the inhibition of their respective RepE sRNA, as *repC^pAt^* and *repC^pTi^*, but not *repAB^pAt^*or *repAB^pTi^*, were significantly less transcribed in GcrA-depleted cells compared to GcrA-repleted cells (Fig. S7A&B). Despite these slight differences, GcrA is apparently one of the key regulatory elements that controls when each extra-element of this complex genome can start its replication so that these replication rounds all finish on time before cell division.

Whether other *Alphaproteobacteria* with RepABC-dependent replicons also use similar GcrA/CcrM-dependent mechanisms of regulation to coordinate their replication with other cell cycle events remains an open question. It is nevertheless relevant to mention that RepABC-dependent replicons are widely distributed among *Alphaproteobacteria* (15), that the GcrA/CcrM couple is found in all the *Alphaproteobacteria* that have multipartite genomes (27,30,48) and that 5’-GANTC-3’ motifs are particularly conserved in *repABC* modules (9). Also, it is specifically in *Alphaproteobacteria* that have more than one essential replicon or essential replication origin, such as *A. tumefaciens* (this study and (16)) or *B. abortus* (32), that the essentially of GcrA has so far been demonstrated, suggesting a potential connection between the essentiality of GcrA and the maintenance of complex genomes in *Alphaproteobacteria*. A connection between DNA methylation and the replication of secondary chromosomes has also been clearly demonstrated in the *Vibrio cholerae Gammaproteobacterium* but it is important to mention that the proteins involved are completely different, engaging the Dam DNA MTase, an RtcB initiator and a secondary chromosomal origin with iterons (10,49,50). Thus, different regulatory mechanisms most likely evolved independently during the domestication of large plasmids that acquired essential genes in different bacterial classes, but their architecture somewhat converged towards epigenetic mechanisms as critical components (51). Further investigations using a diversity of bacteria with complex genomes will now be essential to validate this assumption and explore its impact on genome maintenance.

## Data availability

Metadata and RNASeq data are available in the NCBI BioProject GSE330725 : https://www.ncbi.nlm.nih.gov/geo/query/acc.cgi?acc=GSE330725. Other data that support the findings of this study are available from the corresponding author upon request.

## Supporting information

Volcano plot

Plasmid sequence

Plasmid sequence

Plasmid sequence

## Acknowledgements

We thank Pamela Brown and Xindan Wang for gifts of strains and plasmids. We also thank Jordan Vacheron, Clara Heiman, Julien Luneau and Jessica Burnier for technical help during statistical/bioinformatic/microscopy data collection and analyses. We thank Anna Puzyrko and Corentin Jaboulay for technical assistance. We finally thank Amélie Besombes and Corentin Jaboulay for their feedback on the manuscript and for discussions during the project.

