## Supplementary Information for "Genome replication and cell division control by the methylation-sensitive GcrA regulator in *Agrobacterium tumefaciens*"

**CONTENT:**

Page 2: SUPPLEMENTARY FIGURES WITH LEGENDS

Page 19: SUPPLEMENTARY TABLES WITH CAPTIONS

Page 23: SUPPLEMENTARY METHODS

Page 28: SUPPLEMENTARY REFERENCES

**SUPPLEMENTARY FIGURES WITH LEGENDS:**

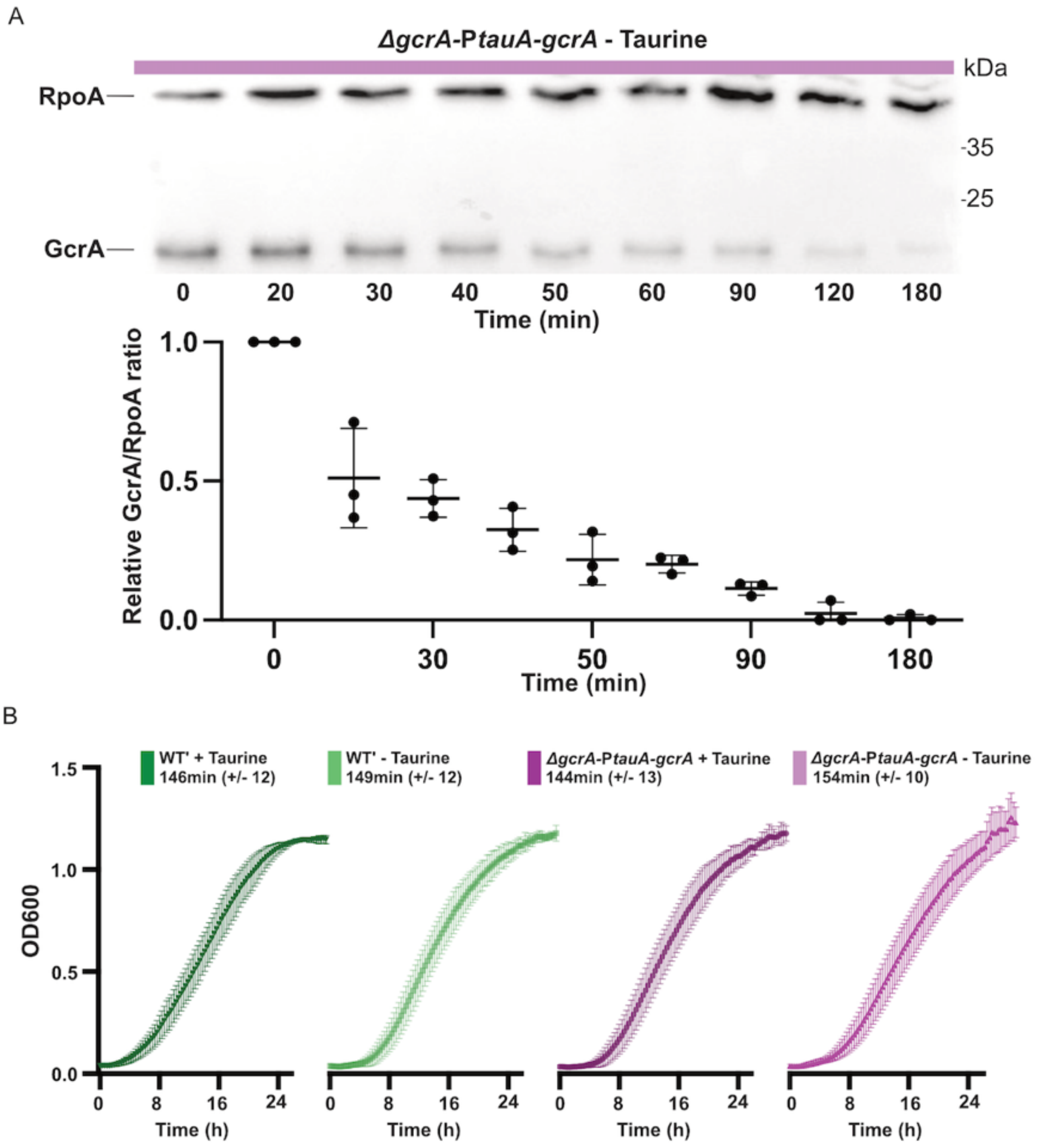


**Figure S1: The half-life of GcrA is much shorter than the generation time of *A. tumefaciens* cells.** (**A**) Estimation of the half-life of GcrA in *A. tumefaciens*. Upper panel: Example of immunoblot showing the rapid decrease of GcrA levels over time in JC2899 (*ΔgcrA PtauA-gcrA*) cells after the removal of taurine from the ATGN medium to switch off *gcrA* transcription. RpoA is used as a loading control. Right side of the images: molecular weight estimations using a prestained protein ladder (PageRuler from Thermo Fisher Scientific, USA). Exponentially growing JC2899 cells cultivated in ATGN + taurine were washed at time 0 and then cultivated in ATGN - taurine. Lower panel: quantification of three independent immunoblots as shown in (A). Band intensities were quantified using the ImageJ software. For each time point, the GcrA/RpoA intensity ratio was calculated on three independent experiments. The GcrA levels/RpoA levels ratios were calculated, and the means were plotted. Comparing the 0 and 15 minute-time points, the half-life of GcrA was estimated as ~15 minutes. (**B**) The growth of JC2141(WT') and JC2899 cultures grown in ATGN +/- taurine was evaluated over-time measuring the $OD_{600nm}$ using a microplate reader (Biotek Synergy H). JC2141 and JC2899 strains were pre-cultured over-night into liquid ATGN + taurine and then washed and resuspended to reach an $OD_{600nm}$~0.1 into the indicated media. Means are plotted and error bars represent standard deviations from three biologically independent replicates. Calculated doubling times (in minutes) for each strain and condition are indicated at the top of each curve.

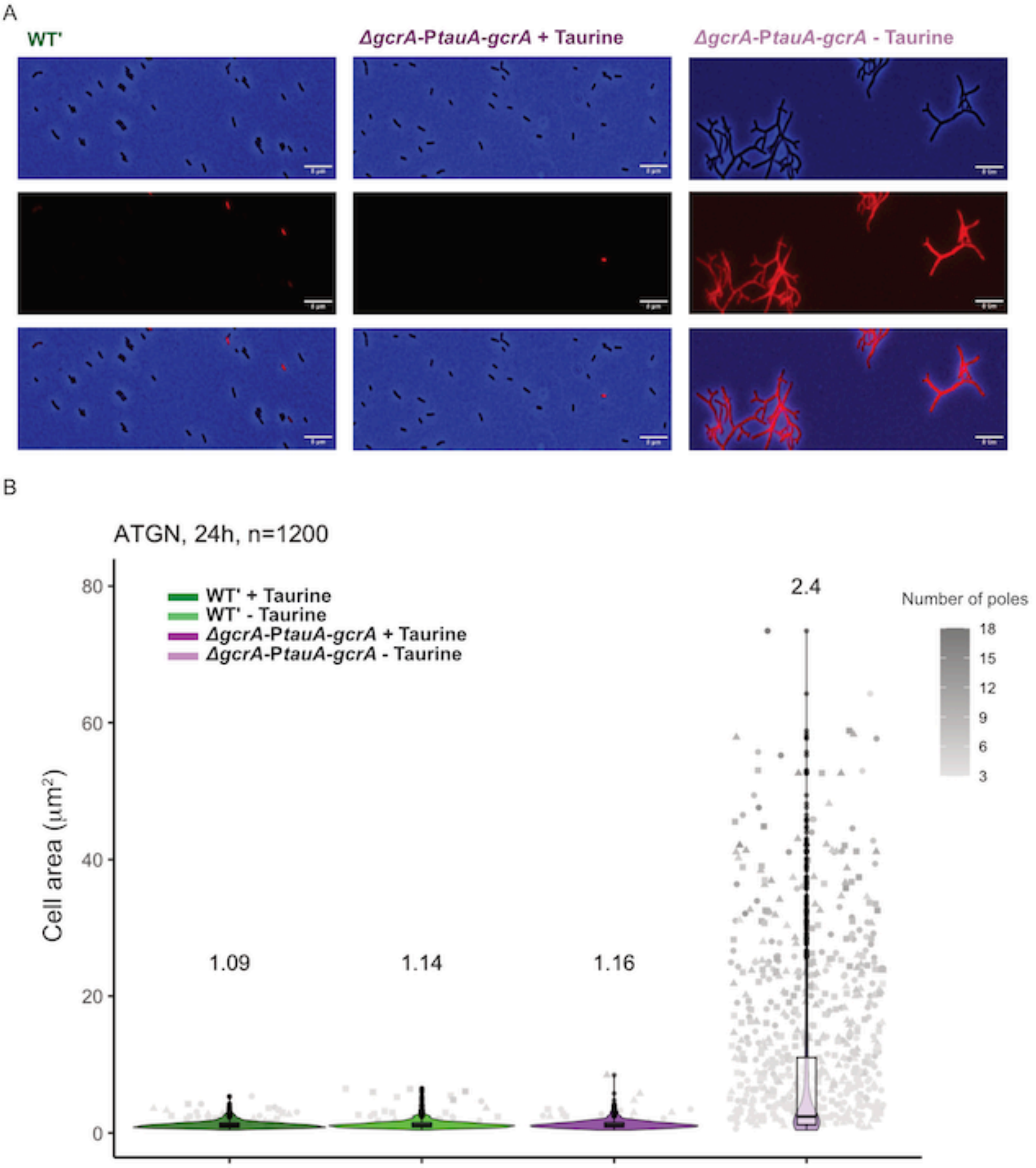


**Figure S2: GcrA-depleted cells elongate, develop multiple poles and die over time.** (A) Live/dead microscopy experiments showing that branching cells are not viable. JC2141 (WT') and JC2899 (*ΔgcrA PtauA-gcrA*) cells were precultured in ATGN + taurine, washed and diluted back in ATGN +/- taurine and cultivated for 24 hours. 50 µl of each culture was stained with the propidium iodide of the LIVE/DEAD *Bac*Light Bacterial Viability Kit (Invitrogen, USA); specifically staining dead cells in red. Cells were then placed on a 1% agarose PBS pad and immediately imaged using the Ph3 (upper image) and DsRED (middle image) filters of the AxioImager M1 microscope (Zeiss); lower images correspond to overlays. These images show that WT' and JC2899 cells cultivated in ATGN + taurine only rarely colored by propidium iodide, while JC2899 cells cultivated in ATGN - taurine are all stained/non-viable. Scale bars correspond to 8 µm. (**B**) Quantification of the number of cell poles/fixed cell cultivated as described above and in Fig. 1D. Three independent replicates were used; 1200 cells were analyzed for each condition. Violin plots of the area of cells are shown for each genotype/condition. The mean of the area of cells is also indicated above each plot. The number of poles of each analyzed cell is also represented by the grey color of dots (triangles and squares correspond to different replicates). Scale bars correspond to 8 µm.

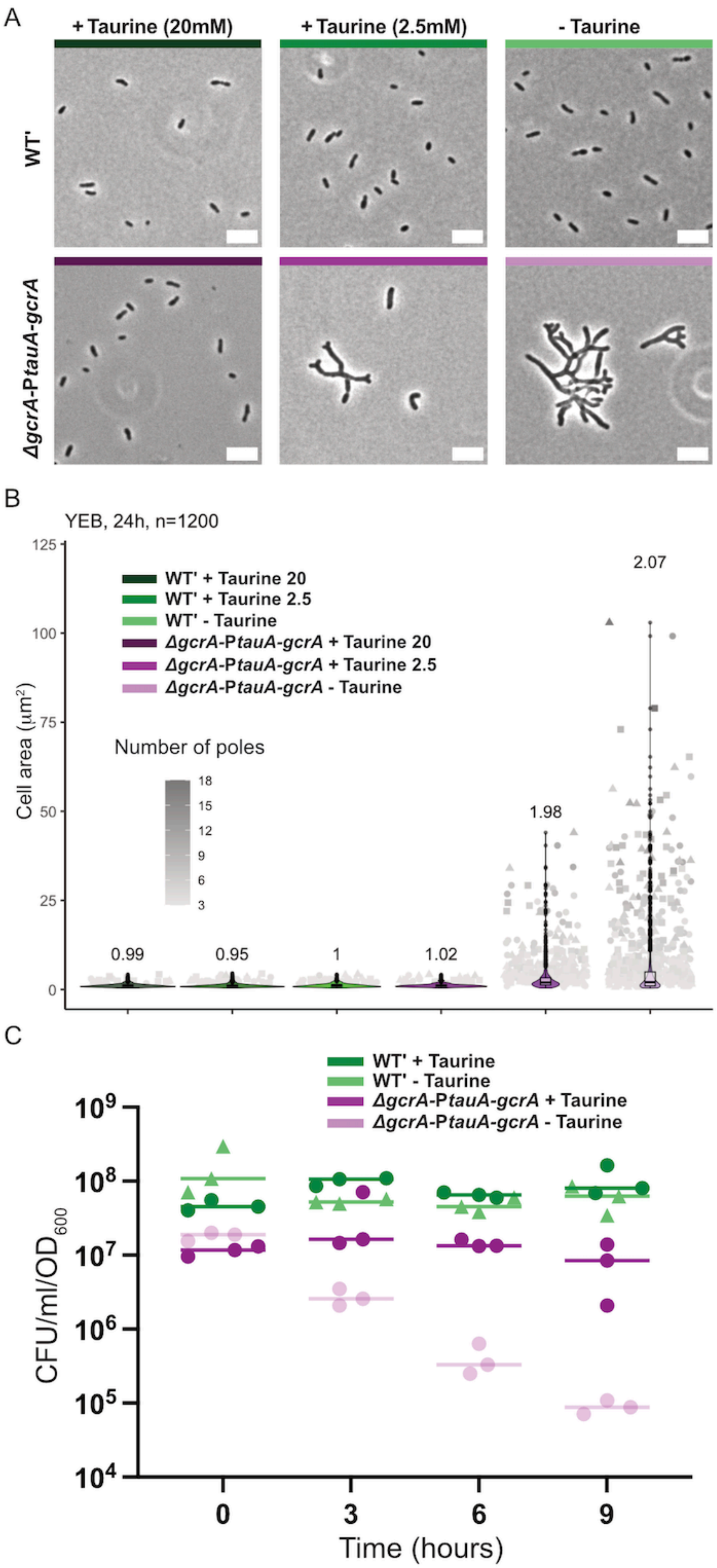
A
+ Taurine (20mM)
+ Taurine (2.5mM)
- Taurine
WT'
ΔgcrA-PtauA-gcrA
B
YEB, 24h, n=1200
Cell area (μm²)
WT' + Taurine 20
WT' + Taurine 2.5
WT' - Taurine
ΔgcrA-PtauA-gcrA + Taurine 20
ΔgcrA-PtauA-gcrA + Taurine 2.5
ΔgcrA-PtauA-gcrA - Taurine
Number of poles
0.99
0.95
1
1.02
1.98
2.07
C
WT' + Taurine
WT' - Taurine
ΔgcrA-PtauA-gcrA + Taurine
ΔgcrA-PtauA-gcrA - Taurine
CFU/ml/OD600
Time (hours)

**Figure S3: Phenotypes of GcrA-depleted cells cultivated in complex medium.** (**A**) Phase-contrast microscopy images of fixed JC2141 (WT') and JC2899 (*ΔgcrA PtauA-gcrA*) cells cultivated in YEB (complex medium) +/- taurine for 24 hours. Scale bars correspond to 5 µm. Taurine was added at 2.5mM or at 20mM as indicated at the top of images. (**B**) Quantification of the number of cell poles/fixed cell cultivated as described above and analyzed and presented as in Fig. S2B. (**C**) Quantification of CFU on ATGNA + taurine from cultures of JC2141 and JC2899 strains in YEB +/- taurine (20mM) for the indicated time. Three independent biological replicates were quantified; horizontal bars correspond to the mean values of the three replicates as done for Fig. 1B.

A

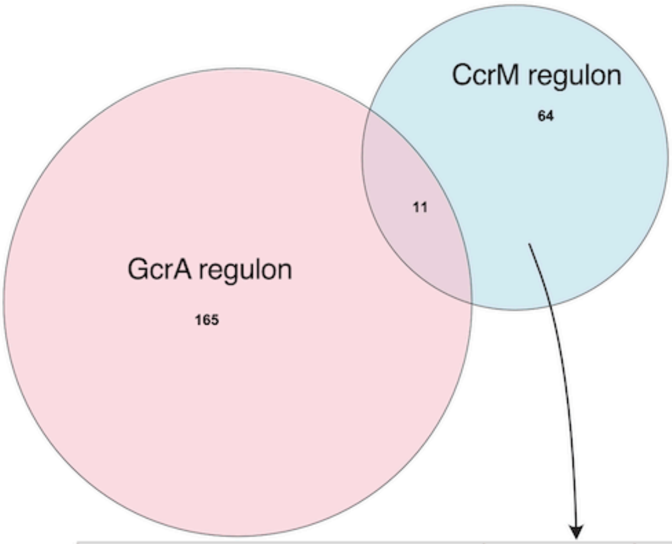


| Protein annotation | *Atu* gene number | GANTC number max. 200 bp upstream ORF | FC GcrA mutant -/+ taurine | FC CcrM mutant -/+ IPTG |
|---|---|---|---|---|
| GcrA | *Atu*0426 | 5 | -123 | +2.8 |
| RepC$^{Ch2}$ | *Atu*3922 | 3 | -6.6 | -5.2 |
| Diguanylate cyclase / GGDEF domain | *Atu*1119 | 1 | -5.4 | -2.1 |
| RepC$^{pTi}$ | *Atu*6045 | 2 | -4.0 | -2.8 |
| Leucyl aminopeptidase | *Atu*0214 | 2 | -3.0 | -2.0 |
| Hypothetical protein | *Atu*0955 | 4 | -2.8 | +2.8 |
| Pyrophosphatase | *Atu*0762 | 1 | -2.7 | +2.4 |
| DgcB / GGDEF domain | *Atu*1691 | 1 | -2.6 | -3.1 |
| TIGR02217 family protein | *Atu*0964 | 1 | -2.5 | +3.3 |
| Phage portal protein | *Atu*0954 | 2 | -2.2 | +2.9 |
| PleC | *Atu*0982 | 5 | +3.0 | +2.9 |

B

| COG categories | COG descriptions |
|---|---|
| C | Energy production and conversion |
| D | Cell cycle control and mitosis |
| E | Amino acid metabolism and transport |
| F | Nucleotide metabolism and transport |
| G | Carbohydrate metabolism and transport |
| H | Coenzyme metabolism |
| I | Lipid metabolism |
| J | Translation |
| K | Transcription |
| L | Replication and repair |
| M | Cell wall/membrane/envelope biogenesis |
| N | Cell motility |
| O | Post-translational modification, protein turnover, chaperones |
| P | norganic ion transport and metabolism |
| Q | Secondary structures |
| R | General function prediction only |
| T | Signal transduction |
| U | Intracellular trafficing and secretion |
| V | Defense mechanisms |
| S | Unknown functions |

**Figure S4: Comparison of the CcrM direct regulon from (1) with the GcrA (direct or indirect) regulon.** (**A**). Upper panel: Venn diagram showing the number of significantly mis-regulated genes (FC>2; adjusted *P*-value <0.01) with at least one 5'-GANTC-3' motif in 200 bp region upstream of each ORF; the CcrM and GcrA regulons are shown in blue and pink, respectively, while the 11 genes belonging to both are shown in violet and listed in the lower panel (the indicated FC corresponds to the fold-change from the RNA-Seq experiments comparing GcrA-depleted with GcrA-repleted cells from Table S4 and CcrM-depleted from CcrM-repleted cells from (1)). (**B**) One-letter code and description of the COGs listed in Fig 2B.

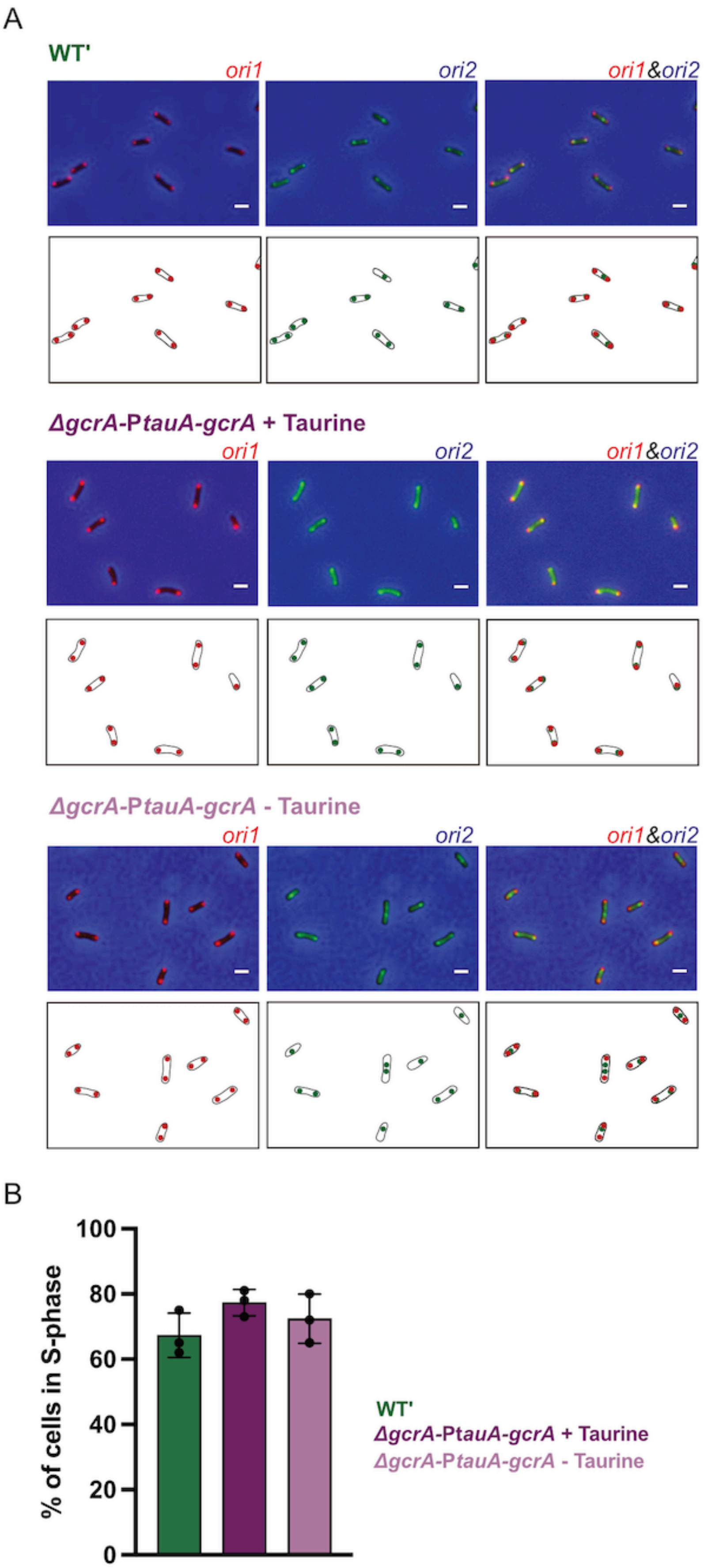
A
WT'
ori1
ori2
ori1&ori2
ΔgcrA-PtauA-gcrA + Taurine
ori1
ori2
ori1&ori2
ΔgcrA-PtauA-gcrA - Taurine
ori1
ori2
ori1&ori2
B
% of cells in S-phase
100
80
60
40
20
0
WT'
ΔgcrA-PtauA-gcrA + Taurine
ΔgcrA-PtauA-gcrA - Taurine

**Figure S5: Localization and number of *ori1* and *ori2* foci per cell depleted for GcrA for 3 hours.** (**A**) Selected microscopy images showing JC3095 (JC2899 with *ori1/mcherry* and *ori2/ygfp* reporters) and JC2777 (JC2141 (WT') with *ori1/mcherry* and *ori2/ygfp* reporters) cells cultured as described in Fig. 3B&C (3 hours +/- taurine). Upper panels for each genotype: overlays of Ph3 and GFP and/or RFP images. Scale bars correspond to 2 µm. Lower panels: schematics showing the subcellular localization of *ori1* (red color) and/or *ori2* (green color) in cells. (**B**) Proportion of S-phase cells in JC3095 and JC2777 populations used for Fig. 3 quantified from images as showed in (A). The number of *ori1* (red foci)/cell was measured (> 1000 cells) to evaluate the % of S-phase cells for each condition/genotype condition (= number of cells with 2 *ori1*/total number of cells). The mean % of S-phase cells from three independent experiments was plotted for each genotype/condition. Error bars correspond to standard deviations.

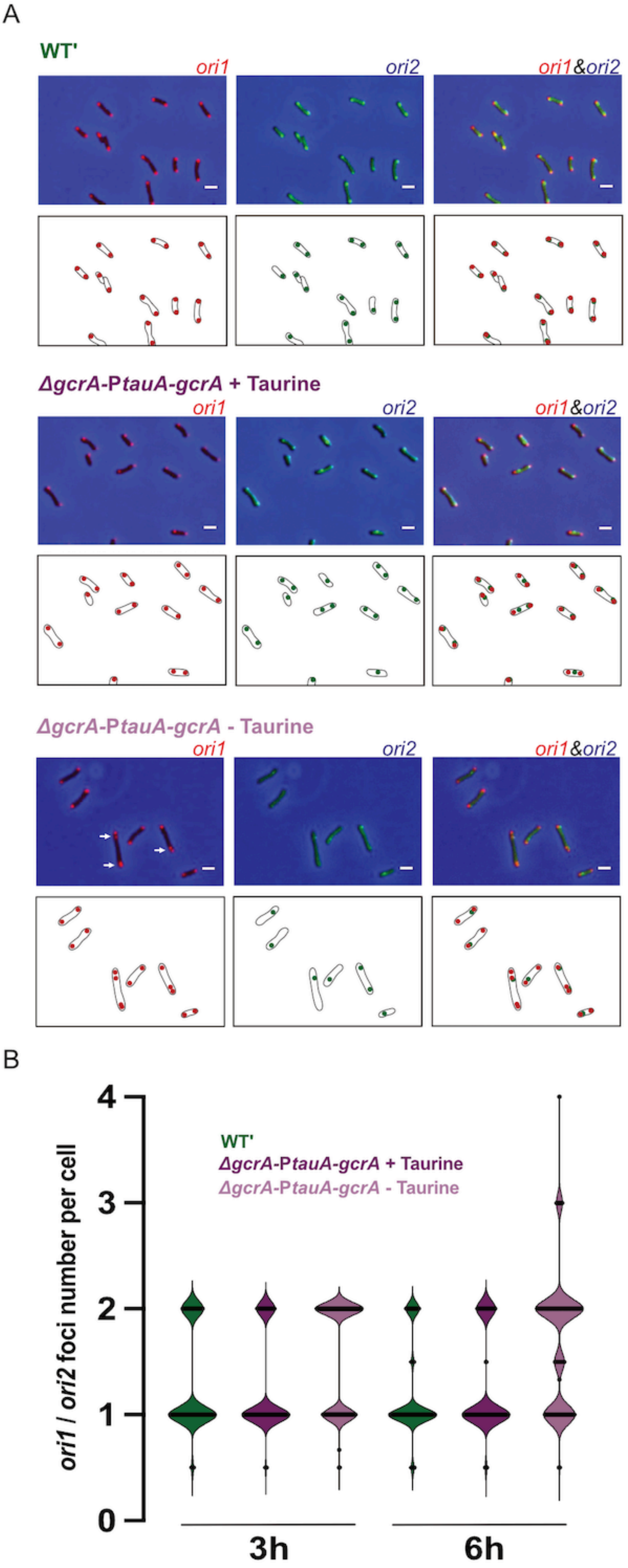

A
WT'
ori1
ori2
ori1&ori2
ΔgcrA-PtauA-gcrA + Taurine
ori1
ori2
ori1&ori2
ΔgcrA-PtauA-gcrA - Taurine
ori1
ori2
ori1&ori2
B
WT'
ΔgcrA-PtauA-gcrA + Taurine
ΔgcrA-PtauA-gcrA - Taurine
ori1 / ori2 foci number per cell
0
1
2
3
4
3h
6h

**Figure S6: Localization and number of *ori1* and *ori2* foci per cell depleted for GcrA for 6 hours.** (**A**) Selected microscopy images showing JC3095 and JC2777 cells cultured, imaged and displayed as Fig. S5A except that cells were cultivated in ATGN +/- taurine for 6 hours instead of 3. White arrows indicate GcrA-depleted cells with more than two *ori1* and only 1 or 2 *ori2*, indicating that initiation from *ori2* can stay blocked even in cells where initiation from *ori1* continues to happen over time. Scale bars correspond to 2 µm. (**B**) Violin plot showing the *ori1*/*ori2* copy number per cell ratio after 3 or 6 hours of growth in ATGN +/- taurine as in Fig. 3D (>1000 cells per condition/genotype). This ratio increases over time in GcrA-depleted cells (JC3095 cultivated ATGN - taurine), indicating that many cells that can initiate at *ori1* cannot initiate at *ori2*.

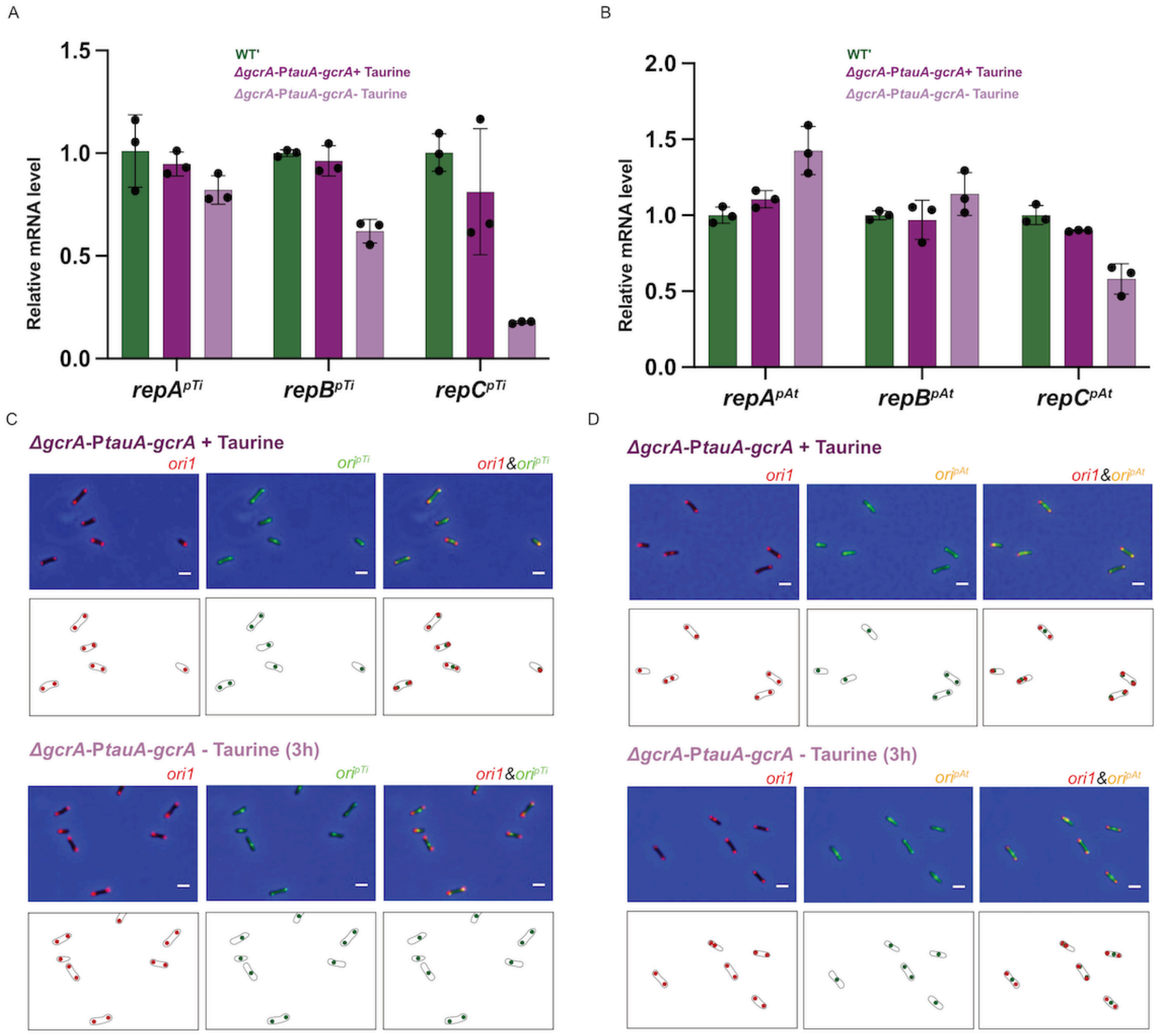


**Figure S7: GcrA affects the expression and the activity of the *repABC*$^{pTi/pAt}$ modules controlling the replication of the pTi and pAt megaplasmids.** (**A**) GcrA is a stronger activator of *repC*$^{pTi}$ than *repBC*$^{pTi}$ expression in *A. tumefaciens*. JC2141 (WT') or JC2899 (*ΔgcrA PtauA-gcrA*) cells were cultivated as in Fig. 2A prior to total RNA extraction for qRT-PCR analyses targeting one ORF at a time. The graph shows relative mRNA levels for each strain/condition. Three biological replicates with three technical replicates were used for each strain/condition. (**B**) GcrA activates *repC*$^{pAt}$ but not *repBC*$^{pAt}$ expression in *A. tumefaciens*. Plots were obtained as for (A). (**C**) and (**D**) GcrA promotes the replication of the pAt and pTi megaplasmids. Selected microscopy images of JC3097 (JC2899 with *ori1/mcherry* and *ori*$^{pTi}$*/ygfp* reporters) and JC3096 (JC2899 with *ori1/mcherry* and *ori*$^{pAt}$*/ygfp* reporters) cells cultured as described in Fig. 3E (3 hours +/- taurine). Upper panels for each genotype/condition: overlays of Ph3 and GFP and/or RFP images. Scale bars correspond to 2 µm. Lower panels: schematics showing the subcellular localization of *ori1* (red color) and/or *ori*$^{pTi}$ or *ori*$^{pAt}$ (green color) in cells.

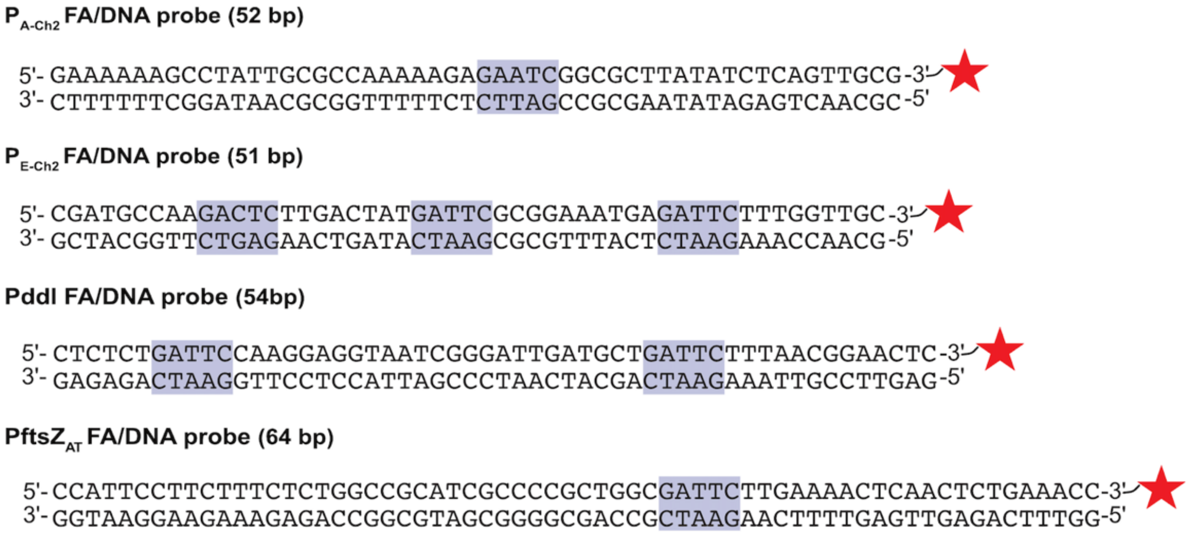


**Figure S8: DNA sequence and 3’-end labeling of the probes used for fluorescence anisotropy (FA) experiments in Figs. 4&6.** 5’-GANTC-3’ motifs are highlighted.

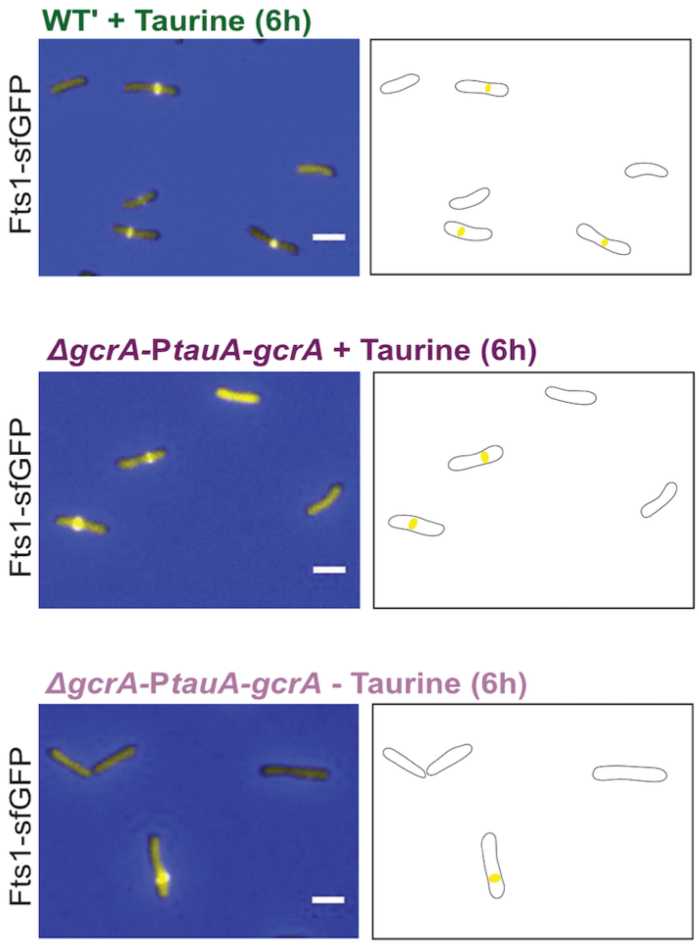


**Figure S9: Impact of GcrA on the assembly of the FtsZ1-sfGFP ring.** Representative images as those used to create Fig. 5C. Strains and growth conditions were the same as described for Fig. 5A, except that cells were cultivated in ATGN +/- taurine for 6 hours. Left panels: overlap of representative Ph3/GFP microscopy images of cells. Right panels: schematics showing the subcellular localization of FtsZ1-sfGFP foci when detectable. Scale bars correspond to 2 µm.

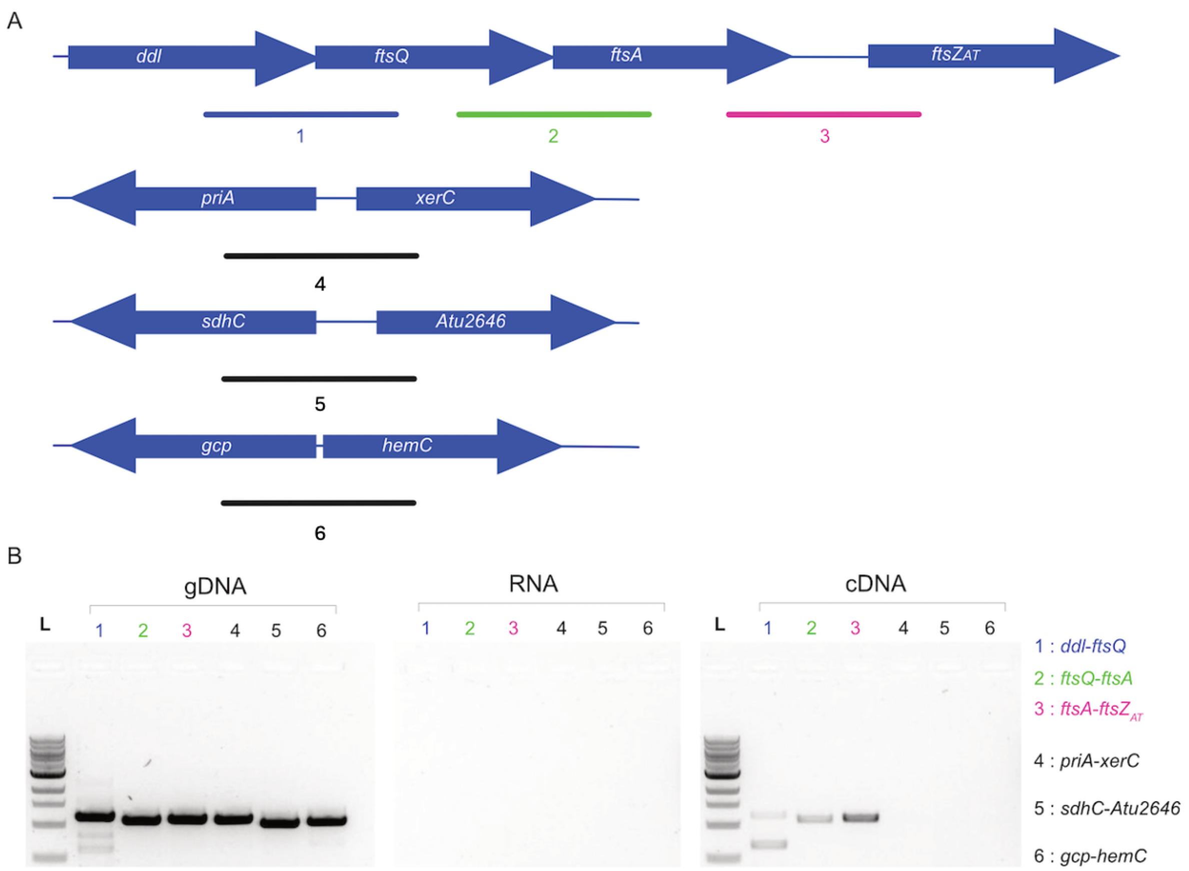


**Figure S10: The *ddl-ftsQAZ$_{AT}$* genes belong to a single operon as they are co-transcribed**

(**A**) Schematic of the *ddl-ftsQAZ$_{AT}$* chromosomal region. Under the schematic, the expected PCR products (1, 2 and 3) that can be detected if gene pairs (*ddl-ftsQ*, *ftsQ-ftsA* and *ftsA-ftsZ$_{AT}$*, respectively) are co-transcribed are depicted. The *priA-xerC*, *sdhC-Atu2646* and *gcp-hemC* genes pairs correspond to three negative controls (PCR 4, 5 and 6) since these genes cannot be co-transcribed as they are in opposite directions. (**B**) Selected agarose gel images showing PCR products obtained using gDNA, total RNA and cDNA from JC2141 cells (cultivated exponentially in ATGN - taurine) as amplification templates and used oligos are listed in Table S3. L: 1 kbp DNA ladder (New England Biolabs, USA).

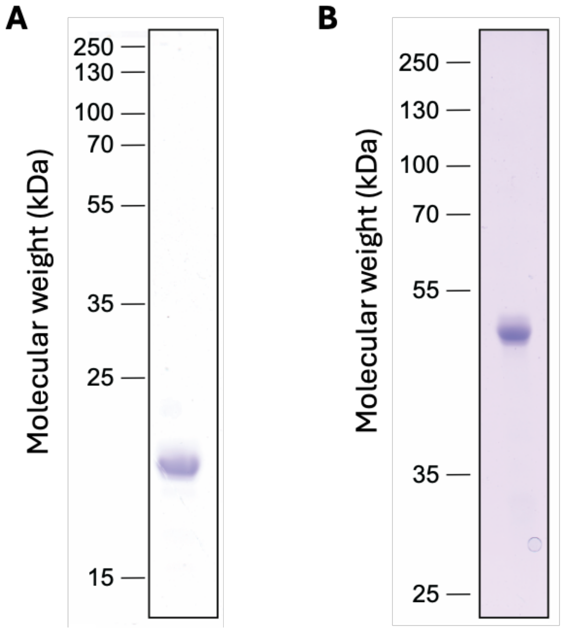


**Figure S11: SDS-PAGE analyzes of purified proteins used in this study.** (**A**) Image of an SDS-PAGE (15%) gel stained with Coomassie Brilliant Blue to check the purity of the His6-GcrA protein (expected molecular weight: ~20 kDa) preparation used to generate antibodies. (**B**) Image of an SDS-PAGE (12%) gel stained with Coomassie Brilliant Blue to check the purity of the GcrA-CDP-His8 protein (expected molecular weight: ~44 kDa) preparation used during fluorescence anisotropy experiments.

### SUPPLEMENTARY TABLES:

#### Table S1: Strains used in this study

| Strain name | Genotype | Description | Reference/Origin |
|---|---|---|---|
| ***Agrobacterium tumefaciens C58 (with one dicentric chromosome)*** | | | |
| JC2141 (WT') | C58 *ΔtetRA::*a-*att*Tn*7* | *tetRA* locus replaced with an artificial *att*Tn*7* site on the dicentric chromosome of *A. tumefaciens* C58 | (1,2) |
| JC2899 | WT' Δ*gcrA* P*tauA-gcrA* | P*tauA-gcrA* inserted at the *ΔtetRA*::a-*att*Tn7 site of the JC2141/WT' chromosome and deletion of the *gcrA* ORF | (1) |
| JC2777 | WT' PT7-*mcherry-parB$^{P1}$- parS$^{P1}$* inserted between *Atu0047* and *Atu0048* and PT7-*ygfp-parB$^{MT1}$-parS$^{MT1}$* inserted between *Atu3973* and *Atu3974* | PT7-*mcherry-parB$^{P1}$-parS$^{P1}$* reporter near *Atu0048* (50kbp away from *ori1*) and PT7-*ygfp-parB$^{pMT1}$-parS$^{pMT1}$* reporter near *Atu3973* (57kbp away from *ori2*) | (1) |
| JC3095 | JC2899 PT7-*mcherry-parB$^{P1}$- parS$^{P1}$* inserted between *Atu0047* and *Atu0048* and PT7-*ygfp-parB$^{MT1}$-parS$^{MT1}$* inserted between *Atu*3973 and *Atu3974* | PT7-*mcherry-parB$^{P1}$-parS$^{P1}$* reporter near *Atu0048* (50kbp away from *ori1*) and PT7-*ygfp-parB$^{pMT1}$-parS$^{pMT1}$* reporter near *Atu3973* (57kbp away from *ori2*) | This study |
| JC3096 | JC2899 PT7-*mcherry-parB$^{P1}$- parS$^{P1}$* inserted between *Atu0047* and *Atu0048* and PT7-*ygfp-parB$^{MT1}$-parS$^{MT1}$* inserted between *Atu6047* and *Atu6048* | PT7-*mcherry-parB$^{P1}$-parS$^{P1}$* reporter near *Atu0048* (50kbp away from *ori1*) and PT7-*ygfp-parB$^{pMT1}$-parS$^{pMT1}$* reporter near Atu6047 (4 kbp away from *ori$^{pTi}$*) | This study |
| JC3097 | JC2899 PT7-*mcherry-parB$^{P1}$- parS$^{P1}$* inserted between *Atu0047* and *Atu0048* and PT7-*ygfp-parB$^{MT1}$-parS$^{MT1}$* inserted between *Atu5336* and *Atu5337* | PT7-*mcherry-parB$^{P1}$-parS$^{P1}$* reporter near *Atu0048* (50kbp away from *ori1*) and PT7-*ygfp-parB$^{pMT1}$-parS$^{pMT1}$* reporter near *Atu5336* (11 kbp away from *ori$^{pAt}$*) | This study |
| ***Escherichia coli*** | | | |
| TOP10 | F- *mcrA* Δ(*mrr-hsdRMS-mcrBC*) φ80l*acZΔM15 ΔlacX74 nupG recA1 araD139 Δ(ara-leu)7697 galE15 galK16 rpsL(Str$^{R}$) endA1 λ$^{-}$* | Used for cloning procedures | Thermo Fisher Scientific (USA) |
| JC1710 | S17.1 λpir | Used for plasmid transfers from *E. coli* to *A. tumefaciens* (conjugation). | (3) |

| | | | |
|---|---|---|---|
| BL21-Gold (DE3) | | Used for protein over-expression from pSG-derived vectors | Sigma-Aldrich (USA) |
| ***Caulobacter crescentus*** | | | |
| JC450 | Synchronizable CB15N wild-type strain (NA1000) | Used for promoter activities measurement | (4) |
| LS3707 | NA1000 *ΔgcrA* P*xyl-gcrA* | Used for promoter activities measurement | (5) |

**Table S2: Plasmids used in this study**

| Plasmid name | Description | Reference/origin |
|---|---|---|
| pOT1e | pBBR1 with promoter-less *egfp,* GentR | (6) |
| $pP_A$(WT)-egfp | pOT1e with a WT $P_{A\text{-}Ch2}$ promoter region (see Fig.4A) controlling *egfp* expression | (1) |
| $pP_E$(WT)-egfp | pOT1e with a WT $P_{E\text{-}Ch2}$ promoter region (see Fig.4A) controlling *egfp* expression | (1) |
| $pP_{ddl\text{-}WT}$ | pOT1e with a WT $P_{ddl}$ promoter region (see Fig.6A) controlling *egfp* expression | This study |
| $pP_{ddl\text{-}\Delta 1}$ | pOT1e with a truncated $P_{ddl}$ promoter region (see Fig.6A) controlling *egfp* expression: 311 bp deletion from the 5' end | This study |
| $pP_{ddl\text{-}\Delta 1M3}$ | pOT1e with a truncated $P_{ddl}$ promoter region (see Fig.6A) controlling *egfp* expression: 311 bp deletion from the 5' end and with the M3 GANTC motif in $P_{ddl\text{-}\Delta 1}$ replaced by a GTNTC motif | This study |
| $pP_{ddl\text{-}\Delta 2}$ | pOT1e with a truncated $P_{ddl}$ promoter region (see Fig.6A) controlling *egfp* expression: 350 bp deletion from the 5' end | This study |
| $pP_{ddl\text{-}\Delta 3}$ | pOT1e with a truncated $P_{ddl}$ promoter region (see Fig.6A) controlling *egfp* expression: 400 bp deletion from the 5' end | This study |
| $pP_{ddl\text{-}\Delta 4}$ | pOT1e with a truncated $P_{ddl}$ promoter region (see Fig.6A) controlling *egfp* expression: 450 bp deletion from the 5' end | This study |
| $pP_{ddl\text{-}\Delta 5}$ | pOT1e with a truncated $P_{ddl}$ promoter region (see Fig.6A) controlling *egfp* expression: 492 bp deletion from the 5' end | This study |
| $pP_{ftsZAT}$ | pOT1e with a WT $P_{ftsZAT}$ promoter region (see Fig.6A) controlling *egfp* expression | This study |
| pWX995 | pNPTS138-derived suicide vector to integrate PT7- *mcherry-parB^P1^-parS^P1^* near *Atu0048* | (7) |
| pWX967 | pNPTS138-derived suicide vector to integrate PT7-*ygfp-parB^MT1^-parS^MT1^* near *Atu3973* | (8) |
| pWX1004 | pNPTS138- derived suicide vector to integrate PT7-*ygfp-parB^MT1-^parS^MT1^* near *Atu6047* | (8) |
| pWX1005 | pNPTS138- derived suicide vector to integrate PT7-*ygfp-parB^MT1^-parS^MT1^* near *Atu5337* | (8) |
| pRV-P*van-ftsZ1-sfgfp* | pRV-MCS2-derivative plasmid to express a fluorescently-tagged FtsZ1-sfGFP protein under P*van (constitutive expression* in *A. tumefaciens*) | (9) |
| pD(Nt)_His6 | 4G-cloning donor for N-terminal His6 tag (https://www.addgene.org/search/catalog/plasmids/?q=223871) | (10) |

| | | |
|---|---|---|
| pD(Ct)_CPD-His8 | 4G-cloning donor for C-terminal CPD-His8 tag (https://www.addgene.org/search/catalog/plasmids/?q=223890) | (10) |
| pSG7522 | Plasmid used as a *gcrA* ORF donor for BsaI-mediated Golden Gate cloning | This study |
| pSG7594 | Expression plasmid with *his6-gcrA* expression under the control of the T7 promoter | This study |
| pSG7593 | Expression plasmid with *gcrA-CPD-his8* expression under the control of the T7 promoter | This study |

**Table S3: Oligonucleotides used in this study**

| Primer name | Sequence (5'→3') | Used for: |
|---|---|---|
| ***Primers used for cloning procedures and strain constructions:*** | | |
| ffjc152 | gtcataagcttggtttgggttccatttcagatg | Construction of $pP_{ddl\text{-}WT}$ |
| ffjc153 | gtcatactagtcgaaccctctgacaccaactc | Construction of $pP_{ddl\text{-}WT}$ |
| ffjc144 | gtcataagcttgagtggcctcgtggcgtaag | Construction of $pP_{ftsZAT}$ |
| ffjc145 | gtcatactagttttaccattccttctttctctgg | Construction of $pP_{ftsZAT}$ |
| MT1 | ggaatattaggtctcaccatgaactggacagacgagcgag | Construction of pSG7522 |
| MT2 | caagcagtgggtctccatccagccttacggcgctcgctaac | Construction of pSG7522 |
| NPJC08 | ccaagctttatgcttgtaaaccgtt | Construction of $pP_{ddl\text{-}\Delta1}$, $pP_{ddl\text{-}\Delta2}$, $pP_{ddl\text{-}\Delta3}$, $pP_{ddl\text{-}\Delta4}$, $pP_{ddl\text{-}\Delta5}$ |
| NPJC10 | cctgcatgttgcaatccgt | Construction of $pP_{ddl\text{-}\Delta1}$ |
| NPJC11 | aggtaatcgggattgatgctga | Construction of $pP_{ddl\text{-}\Delta2}$ |
| NPJC12 | gcggcggttaacgaaagta | Construction of $pP_{ddl\text{-}\Delta3}$ |
| NPJC13 | gatctcaaggatcatgtgcca | Construction of $pP_{ddl\text{-}\Delta4}$ |
| NPJC14 | ggtcggtttggacatggt | Construction of $pP_{ddl\text{-}\Delta5}$ |
| NPJC16 | ggattgatgctgcttctttaacggga | Construction of $pP_{ddl\text{-}\Delta1M3}$ |
| NPJC18 | ggattgatgctgattctttaacggga | Construction of $pP_{ddl\text{-}\Delta1M3}$ |
| ***Primers used for qRT-PCR:*** | | |
| MS110 | CCGCAGGCTAGTCTTTCAGCA | $repA^{pTi}$ |
| MS111 | CGGTCGGATCACATCTCGCAA | $repA^{pTi}$ |
| MS112 | GGCTGCCGGTCGATATCAGAT | $repB^{pTi}$ |
| MS113 | AGGTCAGCGCGGTTAAGATTT | $repB^{pT}$ |
| MS114 | GGTGCCGAACTGGTCGTAT | $repC^{pTi}$ |
| MS115 | GAGCCCGCAATCAACCAGTAT | $repC^{pTi}$ |
| MS118 | CGGTGCAAACGAGACGATTTA | $repA^{pAt}$ |
| MS119 | GCCCGGAATAAGGTCGATACCAT | $repA^{pAt}$ |
| MS120 | GCAGCCAAGGATCTCGGAATT | $repB^{pAt}$ |
| MS121 | GCGCGCTCGATCCAGGTTA | $repB^{pAt}$ |
| MS122b | GGCAAGTCGGCAGACAAATGGA | $repC^{pAt}$ |
| MS123b | CGCAGTTCGTTCTCGGGATAAA | $repC^{pAt}$ |
| ***Primers used for*** **in vitro** ***experiments*** | | |
| ffjc128 | cgatgccaag-N6-Me-dA-ctcttgactatg-N6-Me-dA-ttcgcggaaatgag-N6-Me-dA-ttctttggttgc-6-FAM | $P_{E\text{-}Ch2}$- probe |
| ffjc129 | gcaaccaaag-N6-Me-dA-atctcatttccgcg-N6-Me-dA-atcatagtcaag-N6-Me-dA-gtcttggcatcg | $P_{E\text{-}Ch2}$- probe |
| ffjc130 | cgatgccaagactcttgactatgattcgcggaaatgagattctttggttgc-6-FAM | $P_{E\text{-}Ch2}$- probe |
| ffjc131 | gcaaccaaagaatctcatttccgcgaatcatagtcaagagtcttggcatcg | $P_{E\text{-}Ch2}$- probe |
| ffjc182 | gaaaaaagcctattgcgccaaaaagag-N6-Me-dA-atcggcgcttatatctcagttgcg-6-FAM | $P_{A\text{-}Ch2}$- probe |
| ffjc183 | cgcaactgagatataagcgccg-N6-Me-dA-ttctctttttggcgcaataggcttttttc | $P_{A\text{-}Ch2}$- probe |

| | | |
|---|---|---|
| ffjc184 | gaaaaaagcctattgcgccaaaaagagaatcggcgcttatatctcagttgcg-6-FAM | $P_{A\text{-}Ch2}$- probe |
| ffjc185 | cgcaactgagatataagcgccgattctctttttggcgcaataggcttttttc | $P_{A\text{-}Ch2}$- probe |
| ffjc186 | ctctctg-N6-Me-dA-ttccaaggaggtaatcgggattgatgctg-N6-Me-dA-ttctttaacggaactc-6-FAM | $P_{ddl}$- probe |
| ffjc187 | gagttccgttaaag-N6-Me-dA-atcagcatcaatcccgattacctccttgg-N6-Me-dA-atcagagag | $P_{ddl}$- probe |
| ffjc188 | ctctctgattccaaggaggtaatcgggattgatgctgattctttaacggaactc-6-FAM | $P_{ddl}$- probe |
| ffjc189 | gagttccgttaaagaatcagcatcaatcccgattacctccttggaatcagagag | $P_{ddl}$- probe |
| ffjc138 | ccattccttctttctctggccgcatcgccccgctggccg-N6-Me-dA-ttcttgaaaactcaactctgaaacc-6-FAM | $P_{ftsZAT}$- probe |
| ffjc139 | ggtttcagagttgagttttcaag-N6-Me-dA-atcggccagcggggcgatgcggccagagaaagaaggaatgg | $P_{ftsZAT}$- probe |
| ffjc140 | ccattccttctttctctggccgcatcgccccgctggccgattcttgaaaactcaactctgaaacc-6-FAM | $P_{ftsZAT}$- probe |
| ffjc141 | ggtttcagagttgagttttcaagaatcggccagcggggcgatgcggccagagaaagaaggaatgg | $P_{ftsZAT}$- probe |
| ***Primers used for PCR to detect ddl-ftsQAZ$_{AT}$ cDNA*** | | |
| ffjc158 | cgtactcgatgagctgcgcc | *ddl-ftsQ* PCR |
| ffjc159 | ggaagctggacgagcttgcg | *ddl-ftsQ* PCR |
| ffjc160 | gtcgacaccgagaagatcgc | *ftsQ-ftsA* PCR |
| ffjc162 | cgcaagctcgtccagcttcc | *ftsQ-ftsA* PCR |
| ffjc163 | gcgatcttctcggtgtcgac | *ftsA-ftsZ$_{AT}$* PCR |
| ffjc164 | ccgagaccttcggtgacgttg | *ftsA-ftsZ$_{AT}$* PCR |
| ffjc166 | cagcgacccagtccacgaag | *priA-xerC* PCR |
| ffjc167 | cgtccgcggccagagcatag | *priA-xerC* PCR |
| ffjc168 | cgcggtcagacagatcgacg | *sdhC-Atu2646* PCR |
| ffjc169 | ggcctgatgatgcagacgcc | *sdhC-Atu2646* PCR |
| ffjc170 | ccggagacgagcaacatcag | *gcp-hemC* PCR |
| ffjc171 | gcacgttgtccttgccgagc | *gcp-hemC* PCR |

**Table S4 (available as Supplementary Excel file): RNA-Seq results comparing the transcriptome of JC2899 (*ΔgcrA* P*tauA-gcrA*) cells cultivated in ATGN -/+ taurine (3 hours).** Replicon NC_003062.2 corresponds to Ch1 in non-fusion strains, replicon NC_003063.2 corresponds to Ch2 in non-fusion strains, replicon NC_003064.2 corresponds to pAt and replicon NC_003065.2 corresponds to pTi. Genes directly or indirectly activated or repressed by GcrA (logFC < -1 or > +1) are highlighted in green or orange, respectively. Adjusted P-values > 0.01 (non-significant changes) are highlighted in grey. Genes of particular interest that are discussed in this study are written in red. COG annotations were done as described before (1). GANTC motifs in 200 bp sequences upstream of the annotated start codon of each gene were also searched as described before (1).

### SUPPLEMENTARY METHODS:

#### Bacterial growth conditions

- *E. coli* strains were grown using standard growth conditions in/on Luria-Bertani (LB: 1% tryptone, 0.5% yeast extract, 1% NaCl, pH 7) +/- bacto-agar (1.5%) medium at 37°C. Antibiotics were added at the following concentrations when needed (liquid/solid media): kanamycin 30/50 µg/mL, gentamycin 15/20 µg/mL.

- *A. tumefaciens* strains were grown at 28°C, Yeast Extract Beef (26) (YEB recipe from fisher scientific Bioworld 306270311: 0.5% tryptone, 0.1% yeast extract, 0.5% nutrient broth powder, 0.5% sucrose, 0.049% MgSO4-7H2O, pH 7.2) or AT minimal medium (without exogenous iron) with 0.5% glucose (ATGN) or 5% sucrose (ATSN). Agar plates and soft-agar plates were prepared with 1.35% (ATGN), 1.2% (ATSN) or 0.25% (soft) bacto-agar, respectively. Antibiotics were added at the following concentrations when needed (liquid/solid media): kanamycin 150/300 µg/mL in all media, gentamycin 100/200 µg/mL in ATGN/ATSN. When indicated, Taurine was added at a final concentration of 2.5/20mM (liquid/solid media), respectively.

- *C. crescentus* strains were grown at 28°C in/on peptone yeast extract (PYE) complex medium or M2G minimal medium, supplemented or not with bacto-agar 1.5%. When indicated, gentamycin was added at the following concentrations: 1µg/ml for liquid and 5 µg/ml for solid PYE media. In M2G liquid medium, gentamycin was added at 40 µg/ml. When indicated, 0.03% xylose was added.

#### Construction of plasmids and strains

- *Construction of the JC3095/JC3096/JC3097 strains:* The pWX995 plasmid (donor of *ori1/mcherry* reporter) was introduced into JC2899 with a kanamycin-resistance selection on ATGN + taurine. Resulting transconjugants were cultivated over-night in ATGN + taurine medium (without antibiotics, for pWX995 re-excision step) before plating onto ATSN + taurine medium (sucrose selection). Kanamycin-sensitive and sucrose-resistant colonies were then selected onto ATGN + taurine. We then screened for colonies that kept the *ori1/mcherry* reporter by fluorescence microscopy. The pWX967, pWX1004 and pWX1005 plasmids (donor of *ori2/ygfp*, $ori^{pTi}$*/ygfp* and $ori2^{pAt}$*/ygfp* reporters respectively) was then introduced into this kanamycin-sensitive strain by conjugation using a kanamycin-resistance selection on ATGN + taurine.

- *Construction of the* $pP_{ddl\text{-}WT}$ *plasmid:* The WT 543bp *ddl* promoter region (shown in Fig. 6A) of the *A. tumefaciens* C58 strain was amplified from JC2141 genomic DNA using primers ffjc152/ffjc153, digested by HindIII/SpeI and cloned into HindIII/SpeI-digested pOT1e to create $pP_{ddl\text{-}WT}$ where the expression of *egfp* is under the control of $P_{ddl}$.

- *Construction of the* $pP_{ddl\text{-}\Delta 1}$, $pP_{ddl\text{-}\Delta 1M3}$, $pP_{ddl\text{-}\Delta 2}$, $pP_{ddl\text{-}\Delta 3}$, $pP_{ddl\text{-}\Delta 4}$, *and* $pP_{ddl\text{-}\Delta 5}$ *plasmids:* $P_{ddl}$ deletion mutants were obtained by PCR from $pP_{ddl\text{-}WT}$ using the primers listed in Table S3: NPJC08 was used as the reverse primer and NPJC10, NPJC11, NPJC12, NPJC13, NPJC14 as the forward primers to obtain $pP_{ddl\text{-}\Delta 1}$, $pP_{ddl\text{-}\Delta 2}$, $pP_{ddl\text{-}\Delta 3}$, $pP_{ddl\text{-}\Delta 4}$, $pP_{ddl\text{-}\Delta 5}$, respectively. The $pP_{ddl\text{-}\Delta 1M3}$ plasmid was obtained by site-directed mutagenesis PCR using primers NPJC16/NPJC18 and $pP_{ddl\text{-}WT}$ as the DNA template.

- *Construction of* $pP_{ftsZAT}$*:* The 93bp region upstream of the *ftsZ*$_{AT}$ ORF (see Fig. 6A) on the *A. tumefaciens* C58 chromosome was amplified from JC2141 genomic DNA using primers ffjc144/ffjc145, and then digested and cloned into pOT1e as described above for $pP_{ddl\text{-}WT}$.

- *Construction of pSG7522, pSG7594 and pSG7593*: A *gcrA* ORF donor for BsaI-mediated Golden Gate cloning (pSG7522) was created by PCR using primers MT1 and MT2 on JC2141 genomic DNA, followed by Gibson Assembly into a linearized pD vector as described in (10), giving pSG7522 (complete sequence as Supplementary .dna file). Donors for N-terminal His6 (pD(Nt)_His6) or C-terminal Cysteine Protease Domain (CPD)-His8 (pD(Ct)_CPD-His8) tags were described in (10). The two pET-based bacterial expression vectors were constructed by inserting a BsaI-linearization module between the T7 promoter and the T7 terminator, so that the BsaI-generated overhangs allowed the insertion of the *gcrA* ORF in frame with the N-terminal or the C-terminal tag by BsaI-mediated Golden-Gate cloning. This step was carried out in a PCR machine (30 cycles of 2 min. at 37°C / 5 min. at 16°C, followed by 10 min. at 50°C) and the 10 µl reaction sample was prepared as described in (10). This resulted in vectors pSG7594 and pSG7593 (complete sequences as Supplementary .dna files).

**Checking and quantifying RNA samples for qRT-PCR and RNA-Seq analyses**

The quality of RNA samples was verified: (i) for the absence of DNA contaminations using standard PCR, (ii) on an agarose gel (1% agarose in TAE 1X) and (iii) using a Fragment Analyzer (Agilent Technologies). The RNAs had RQNs between 9.3 and 9.9. Concentrations of RNA samples were measured using a Nanodrop spectrophotometer.

**RNA-Seq experiments and analyses**

RNA-seq libraries were prepared from 100 ng of total RNA with the Illumina Stranded mRNA Prep reagents (Illumina) using a unique dual indexing strategy, and following the official protocol. The polyA selection step was replaced by an rRNA depletion step with the RiboCop for Bacteria, mixed bacterial samples reagents (Lexogen). Libraries were quantified by a fluorometric method (QubIT, Life Technologies) and their quality assessed on a Fragment Analyzer. Sequencing was performed on an Illumina NovaSeq 6000 for 100 cycles single read. Sequencing data were demultiplexed using the bcl2fastq2 Conversion Software (version 2.20, Illumina).

Sequences matching to ribosomal RNA sequences were removed with fastq_screen (v. 0.11.1) (11). Remaining reads were further filtered for low complexity with reaper (v. 15–065) (12).

Reads were aligned against the *Agrobacterium fabrum* C58 ASM9202v1 genome using STAR (v. 2.5.3a) (13). The number of read counts per gene locus was summarized with htseq-count (v. 0.9.1) (14) using a custom *Agrobacterium fabrum* C58 ASM9202v1 gene annotation. Quality of the RNA-seq data alignment was assessed using RSeQC (v. 2.3.7) (15). Counts per gene table was used for statistical analysis in R (R version 4.2.1). Genes with low counts were filtered out according to the rule of 1 count per million (cpm) in at least 3 samples. Library sizes were scaled using TMM normalization (EdgeR package version 3.34.1 from (16)) and log-transformed with limma cpm function (Limma package version 3.54.0 from (17)). Differential expression was computed with limma by fitting the samples into a linear model and performing comparisons with moderated t-tests. Global p-value adjustment with Bonferroni–Hotchberg method was used for all comparisons. Genes with an adjusted P-value < 0.01 and a minimum fold-change of 2 were considered as significantly mis-regulated. Volcano plots showing differentially expressed genes were made in R (v4.2.1) by plotting the log2(fold change) against the –log10(adjusted P-value) for this comparison.

**Expression and purification of His6-GcrA for antibody generation**

The pSG7594 expression vector was introduced into the *E. coli* BL21-Gold (DE3) strain and the resulting strain was inoculated into 1 liter of Terrific Broth (TB) in a non-baffled glass flask. The culture was first grown at 37°C to an $OD_{600nm}$ of ~1 and the culture temperature was then reduced to 20°C. Expression was then induced by the addition of 0.4 mM IPTG and allowed to proceed overnight (16 h). Cells were harvested by centrifugation, resuspended in 70 ml of PBS supplemented with 25 mM Imidazole, and the suspension was then sonicated on ice to lyse cells (Bandelin sonicator, VS70T tip, 10 min. total sonication time with pulses of 1 sec. ON / 1 sec. OFF). The lysate was then clarified by centrifugation at 40000g for 30 min. at 4°C, and the supernatant was loaded onto a 5 ml HisTrap-HP column (Cytiva). After washing the column with 8 column volumes (cV) of PBS, the bound material was eluted with a gradient from 25 mM to 500 mM Imidazole in PBS. Fractions containing His6-GcrA were identified by SDS-PAGE and the purified protein sample (Fig. S11A) was prepared for injection into rabbits following the requirements from Eurogentec (Belgium).

**Expression and purification of GcrA-CPD-His8 protein for Fluorescence Anisotropy experiments**

The pSG7593 expression vector was introduced into the *E. coli* BL21- Gold (DE3) strain and the resulting strain was cultivated as described above for cells containing the pSG7594 vector. Cells were harvested by centrifugation and the cell pellet was resuspended into a lysis buffer (50 mM Tris-HCl pH 7.5, 1000 mM NaCl, 5 % glycerol, 25 mM Imidazole) supplemented with 5 mM beta-mercaptoethanol and protease inhibitors. Sonication and lysate clarification were also performed as described above. The lysate was then loaded onto a 5 ml HisTrap-HP column (Cytiva), which was then washed with 8 column volumes of buffer A (50 mM Tris-HCl pH 7.5, 250 mM NaCl, 5 % glycerol, 25 mM Imidazole, 5 mM beta-mercaptoethanol) and eluted with a 10 cV gradient from buffer A to buffer B (50 mM Tris-HCl pH 7.5, 250 mM NaCl, 5 % glycerol, 500 mM Imidazole, 5 mM beta-mercaptoethanol). Fractions containing GcrA-CPD-His8 target were pooled, loaded onto a 5 ml HiTrap Heparin column (Cytiva), and bound proteins were eluted with a 10 cV gradient from buffer A to buffer C (50 mM Tris-HCl pH 7.5, 1000 mM NaCl, 5 % glycerol, 5 mM beta-mercaptoethanol). Fractions containing sufficiently pure GcrA-CPD-His8 were pooled and diluted to a final protein concentration of 100 μM 50 μl aliquots were snap-frozen and stored at -70°C. The purity of the sample can be seen in Fig. S11B.
